# Adult consequences of repeated nicotine vapor inhalation in adolescent rats

**DOI:** 10.1101/2022.11.17.516984

**Authors:** Arnold Gutierrez, Jacques D. Nguyen, Kevin M. Creehan, Yanabel Grant, Michael A. Taffe

**Affiliations:** Department of Neuroscience; The Scripps Research Institute; La Jolla, CA, USA; Department of Psychiatry, University of California, San Diego; La Jolla, CA, USA; Department of Psychology and Neuroscience, Baylor University; Waco, TX USA

**Author notes:** Address Correspondence to: Dr. Michael A. Taffe, Department of Psychiatry, 9500 Gilman Drive; University of California, San Diego, La Jolla, CA 92093; USA.

## Abstract

**Introduction:** There has been a recent resurgence in nicotine inhalation in adolescents due to the popularity and availability of Electronic Nicotine Delivery Systems (ENDS). Almost five times as many US high-school seniors inhale nicotine vapor daily compared with those who smoke tobacco. This study was conducted to determine the impact of repeated adolescent vapor inhalation of nicotine on behavior in adulthood.

**Methods:** Male and female Sprague-Dawley rats were exposed to 30-minute sessions of ENDS vapor inhalation, twice daily, from Post-Natal Day (PND) 31 to PND 40. Conditions included vapor from the propylene glycol (PG) vehicle or nicotine (30 mg/mL in the PG). Animals were assessed for effects of nicotine on open field (PND 74-105) and wheel activity (PND 126-180) and for the self-administration of nicotine vapor (PND 285-395). Plasma levels of nicotine and cotinine were assessed in separate groups of male and female Wistar and Sprague-Dawley rats after a single nicotine inhalation session.

**Results:** Group mean plasma nicotine ranged from 39 to 59 ng/mL post-session with minimal strain differences detected. Adolescent nicotine exposure modestly enhanced sensitivity to the locomotor stimulating effects of nicotine (0.1-0.8 mg/kg, s.c.) in an open field in female rats, but didn’t change effects of nicotine on wheel activity. Female rats exposed to nicotine (30 mg/mL) vapor as adolescents responded more vigorously than PG exposed females for nicotine vapor in a FR5 challenge.

**Conclusions:** Repeated adolescent nicotine vapor inhalation leads to enhanced liability for nicotine self-administration in adulthood in female rats, but minimal change in spontaneous locomotor behavior.

## Introduction

Adolescent nicotine inhalation surged recently due to the popularity of Electronic Nicotine Delivery Systems (ENDS, “e-cigarettes”). About 11.7% of US twelfth graders, and 6.9% of tenth graders, vaped nicotine daily in 2019, while daily tobacco smoking reached a decades’ low of 2.4% and 1.3%, respectively (Johnston et al., 2021). About 3.3% of middle school students and 14.1% of high school students used ENDS, while only 1.0% and 2.0%, respectively, smoked cigarettes (Park-Lee et al., 2022). About 20-21% of adults ages 18-22 vaped nicotine monthly, compared with 10-13% of those in their later twenties (Schulenberg et al., 2021), showing that adolescent vaping increased sustained use patterns in adulthood.

Research (Fowler et al., 2018) on the health impacts of ENDS exposure to nicotine can be greatly advanced by laboratory models to better isolate risks specific to ENDS from social or environmental factors in human behavior. The goal of this study was to determine lasting consequences of repeated inhalation of nicotine in adolescent rats, using a system proven effective for delivering active doses to adult rats of either sex (Javadi-Paydar et al., 2019b; Lallai et al., 2021; Montanari et al., 2020), to pregnant rats (Breit et al., 2022) and to mice (Cooper, Akers and Henderson, 2021; Echeveste Sanchez et al., 2022; Henderson and Cooper, 2021). ENDS-based systems have shown efficacy for nicotine self-administration in rats (Smith et al., 2020) and mice (Cooper, Akers and Henderson, 2021; Henderson and Cooper, 2021) and repeated daily exposure to nicotine via vapor inhalation leads to withdrawal after discontinuation in rats (Montanari et al., 2020).

Adolescent nicotine *injection* produces lasting effects, as male rats injected with nicotine (1.0 mg/kg, s.c.) daily from Post-Natal Day (PND) 28-41 self-administered more nicotine (Renda et al., 2020), and male rats injected with nicotine, i.v., from PND 28-31 self-administered more methamphetamine (Cardenas and Lotfipour, 2022). Daily nicotine injection (0.6 mg/kg, s.c.) from PND 28-42 enhanced pain responses and reduced the efficacy of morphine when rats were assessed as adults (Khalouzadeh, Azizi and Semnanian, 2022). Relatedly, nicotine inhalation by pregnant dams impairs motor coordination in offspring during adolescence (Hussain, Breit and Thomas, 2022). Perplexingly, adolescent nicotine produced either increased *or* decreased nicotine self-administration in female rats in two similar studies from the same laboratory (Chellian et al., 2021; Chellian et al., 2020). Given that inhalation of drugs can produce a similar subjective high at a lower plasma concentration compared with *intravenous* delivery (Cook et al., 1993), it is critical to determine whether nicotine *inhalation* produces lasting consequences.

Nicotine vapor inhalation models are still relatively new with few data available regarding behavior, as well as plasma levels of nicotine or cotinine, across rat age, sex and strain. One recent report found that Wistar rats exposed to daily nicotine vapor from PND 33-46 exhibited more nosepokes in a nicotine vapor delivery port during passive drug delivery than did air-exposed control groups (Espinoza et al., 2022). Small differences in vapor exposure methods related to vapor puffing protocols, session duration, and rat strain have varied across an emerging literature (Espinoza et al., 2022; Javadi-Paydar et al., 2019b; Montanari et al., 2020) so we first determined plasma concentrations of nicotine and cotinine after exposure in our model. Behavioral and physiological effects of vapor inhalation of nicotine have not yet been reported for adolescent rats, so we next determined if locomotor activity and thermoregulation effects in adult animals (Javadi-Paydar et al., 2019b) generalize to adolescents of each sex. The study then exposed groups of adolescent rats to repeated inhalation of nicotine vapor or that of the vehicle, and assessed them during adulthood for responses to nicotine in spontaneous locomotor behavior in open field and activity wheels, and for nicotine vapor self-administration.

## Methods

### Subjects

Male and (N=22) and female (N=54) Sprague-Dawley (Envigo/Harlan) and male (N=6) and female (N=6) Wistar (Charles River) rats were used for this study. The vivarium was kept on a 12:12 hour reversed light-dark cycle (lights out at 0800); studies were conducted during vivarium dark. Food and water were provided ad libitum in the home cage. Procedures were conducted in accordance with protocols approved by the IACUCs of The Scripps Research Institute and of the University of California, San Diego and were consistent with the NIH Guide (Garber et al., 2011).

### Drugs

Nicotine bitartrate (Sigma-Aldrich, St. Louis, MO) was dissolved in propylene glycol (PG; Fisher Scientific) at a concentration of 30 and 60 mg of nicotine bitartrate per mL of PG for vapor and dissolved in physiological saline for injection (0.2-1.0 mg/kg, s.c.) studies. PG was used as the vehicle for comparability with our prior reports (Javadi-Paydar et al., 2019a; Javadi-Paydar et al., 2019b).

### Plasma Nicotine and Cotinine Analysis

Nicotine and cotinine concentrations were quantified using liquid chromatography/mass spectrometry (LCMS) as previously reported (Javadi-Paydar et al., 2019b). See **Supplemental Materials** for full description.

### Apparatus

#### Vapor Inhalation

A vapor inhalation system (La Jolla Alcohol Research, Inc) previously shown to deliver Δ9-tetrahydrocannabinol (THC), cannabidiol, heroin, oxycodone, methamphetamine and nicotine (Gutierrez, Creehan and Taffe, 2021; Javadi-Paydar et al., 2019a; Javadi-Paydar et al., 2019b; Nguyen et al., 2016; Nguyen et al., 2019) was used. In brief, vapor was delivered into sealed vapor exposure chambers through controllers which trigger commercial e-cigarette tanks/atomizers. Vacuum control through an exhaust flowed room air at ∼1 L per minute and ensured that vapor entered the chamber. See **Supplemental Materials** for full description.

#### Open Field

Locomotor activity was recorded videographically in plastic chambers (80 cm L X 44 cm W X 33 cm H) and analyzed off-line using ANY-maze tracking software (Stoelting). See **Supplemental Materials**.

#### Activity Wheels

Activity on wheels (Med Associates; Model ENV-046) was assessed in 30-minute sessions, as previously described (Gilpin et al., 2011; Miller et al., 2013; Taffe et al., 2021). See **Supplemental Materials** for full description.

### Experiments

#### Experiment 1: Plasma nicotine and cotinine after vapor inhalation or injection

Groups (N=6) of young adult male and female Sprague-Dawley and Wistar rats were implanted with jugular catheters as previously described (Nguyen et al., 2020; Nguyen, Grant and Taffe, 2021; Nguyen et al., 2018). Two common laboratory strains were contrasted to determine if major strain differences might occur, useful for generalization of this relatively new procedure. Experiments were conducted in which blood was collected after a 30-minute nicotine (30 mg/mL) vapor inhalation session (35, 50, 120, 240 minutes after vapor initiation) or after a 1 mg/kg, s.c. injection (15, 30, 60, 120 minutes post-injection). Occasionally a sample could not be obtained, resulting in the sample size as described in the relevant figures. Blood sampling studies were conducted consistent with recommendations for good practices (Diehl et al., 2001), which in part dictated the use of young adults, as intervals for blood volume recovery between experiments would cross the adolescent range.

#### Experiment 2: Effect of repeated adolescent nicotine inhalation on open field locomotion

Groups (N=8 per treatment) of male and female Sprague-Dawley rats arrived in the laboratory at PND 25 and were exposed to either PG or nicotine (30 mg/mL in the PG) vapor in 30-minute sessions twice per day (0800/1400) for 10 days (PND 31 to PND 40). Exposure parameters previously used in adult rats (Javadi-Paydar et al., 2019b) were validated in separate groups of adolescents (See **Supplemental Materials Figure S2**). Exposure was during the vivarium dark cycle but under white light illumination. Locomotor activity was assessed from PND 74 to PND 105 in open field; assessment was conducted for 30-minutes during the vivarium dark cycle but under white light illumination to facilitate video tracking. One 30-minute baseline recording session was conducted to habituate the animals to the procedure and apparatus. Thereafter, the doses 0.0, 0.2, 0.8 mg/kg nicotine, s.c., were assessed in a counter-balanced order and next the 0.0, 0.1, 0.4 mg/kg nicotine, s.c., doses were assessed in a counter-balanced order, a minimum of 3-4 days apart. Injections were 15 minutes prior to evaluation.

#### Experiment 3: Effect of repeated adolescent nicotine inhalation on wheel activity

The Experiment 2 groups were assessed for wheel activity with quarter rotations recorded every 15 minutes during 1 h sessions; these were conducted under white light illumination for better comparison with the open field study. Rats completed two baseline sessions (5-7 days apart, starting PND 126) without pre-treatment for habituation. Thereafter (PND 165-PND180) sessions were run immediately after pre-injection with nicotine (0.0, 0.2, 0.4, 0.8 mg/kg, s.c.) with doses administered 15 minutes prior in a counter-balanced order, with a minimum 3-4 day interval between doses.

#### Experiment 4: Effect of repeated adolescent nicotine inhalation on nicotine self-administration

From PND 285 onward the rats from Experiments 2-3 were assessed for self-administration of nicotine (30 mg/mL) vapor in 30-minute sessions. Four chambers with nose-poke manipulanda and two chambers with lever manipulanda were used, with the assignment counterbalanced across groups. No consistent differences associated with manipulanda were noted throughout the study. A response on the drug-associated manipulandum (FR-1 response requirement) resulted in illumination of the cue light and delivery of a 1 second puff of vapor. This was followed by a 20 second time-out during which the cue light remained illuminated; responses were recorded but led to no consequences. Intervals between successive sessions were [2, 3, 2, 15, 6, 5, 2, 4 days] for the initial 9 acquisition sessions. Animals were next evaluated with a range of concentrations (0, 10, 30 or 60 mg/mL nicotine in the PG) evaluated once per concentration in a counterbalanced order across sequential test sessions, under an FR1 schedule.

Rats were then returned to FR1 / Nicotine (30 mg/mL) for one session, after which the response contingency was increased to FR5. The rationale is that increased responding on the drug-associated manipulandum is interpreted as evidence of drug-seeking, as in (Chaudhri et al., 2005). A facilities flooding emergency disrupted access to testing rooms after the 3rd FR5 session. Animals were idled for three weeks and then re-started under the same FR5 contingency before returning to FR1 for four additional sessions. This latter study was based on a prior report that female rats initiated at 6 weeks of age increased their nicotine intravenous self-administration (IVSA) more than female rats initiated at 8 weeks of age when the contingency was returned to FR1 from FR8 (Levin et al., 2011). Analysis thus focused on the first three FR5 sessions compared with the immediate prior FR1 session. The second analysis compared the post-interruption FR5 sessions and the subsequent FR1 sessions with the FR1 session conducted prior to the FR5 manipulation.

### Data Analysis

Plasma nicotine and cotinine were analyzed by mixed-effect models to account for missing values. A within-subject factor of Time after vapor initiation / injection and a factor for Group were included. Open Field distance in baseline, Open Field distance after nicotine, wheel rotations and vapor deliveries obtained were analyzed by 3-way ANOVA including between-subjects factors for Sex and Adolescent Treatment and a within subject factor for Time, Dose or Session, respectively. Follow-up analyses with 2-factor ANOVA were used where relevant to collapse across non-significant factors and facilitate post-hoc exploration.

A criterion of P<0.05 was used to infer a significant difference. Significant main effects were followed with post-hoc analysis using Tukey (multi-level factors), Sidak (two-level factors) or Dunnett (treatments versus control within group) correction. All analysis used Prism for Windows (v. 9.4.0; GraphPad Software, Inc, San Diego CA).

## Results

### Experiment 1: Plasma nicotine and cotinine

#### Plasma nicotine and cotinine after vapor inhalation

Concentrations of nicotine and cotinine were analyzed across the four Groups (**Figure 1**), and mixed-effects analyses confirmed significant effects of Time, of Group and / or the interaction on plasma *<u>nicotine</u>* [Group: n.s.; Time: F (1.291, 17.21) = 172.4, P<0.0001; Interaction: F (9, 40) = 2.39, P<0.05], and plasma *<u>cotinine</u>* [Group: F (3, 17) = 9.03, P<0.001; Time: F (1.219, 16.25) = 240.4, P<0.0001; Interaction: F (9, 40) = 2.55, P<0.05], concentrations. Follow-up analysis was conducted collapsing first across rat strain [Sex: F (1, 19) = 4.51, P<0.05; Time: F (1.394, 21.37) = 180.7, P<0.0001; Interaction: F (3, 46) = 5.97, P<0.005] and next across rat sex [Strain: n.s.; Time: F (1.247, 19.12) = 141.2, P<0.0001; Interaction: n.s.]. The post-hoc test failed to confirm significant differences between male and female groups at any time-point. Post-hoc analysis of Time, collapsed across group, confirmed that plasma concentrations of nicotine at each time-point were significantly different from the concentrations at every other timepoint.

**Figure 1:**
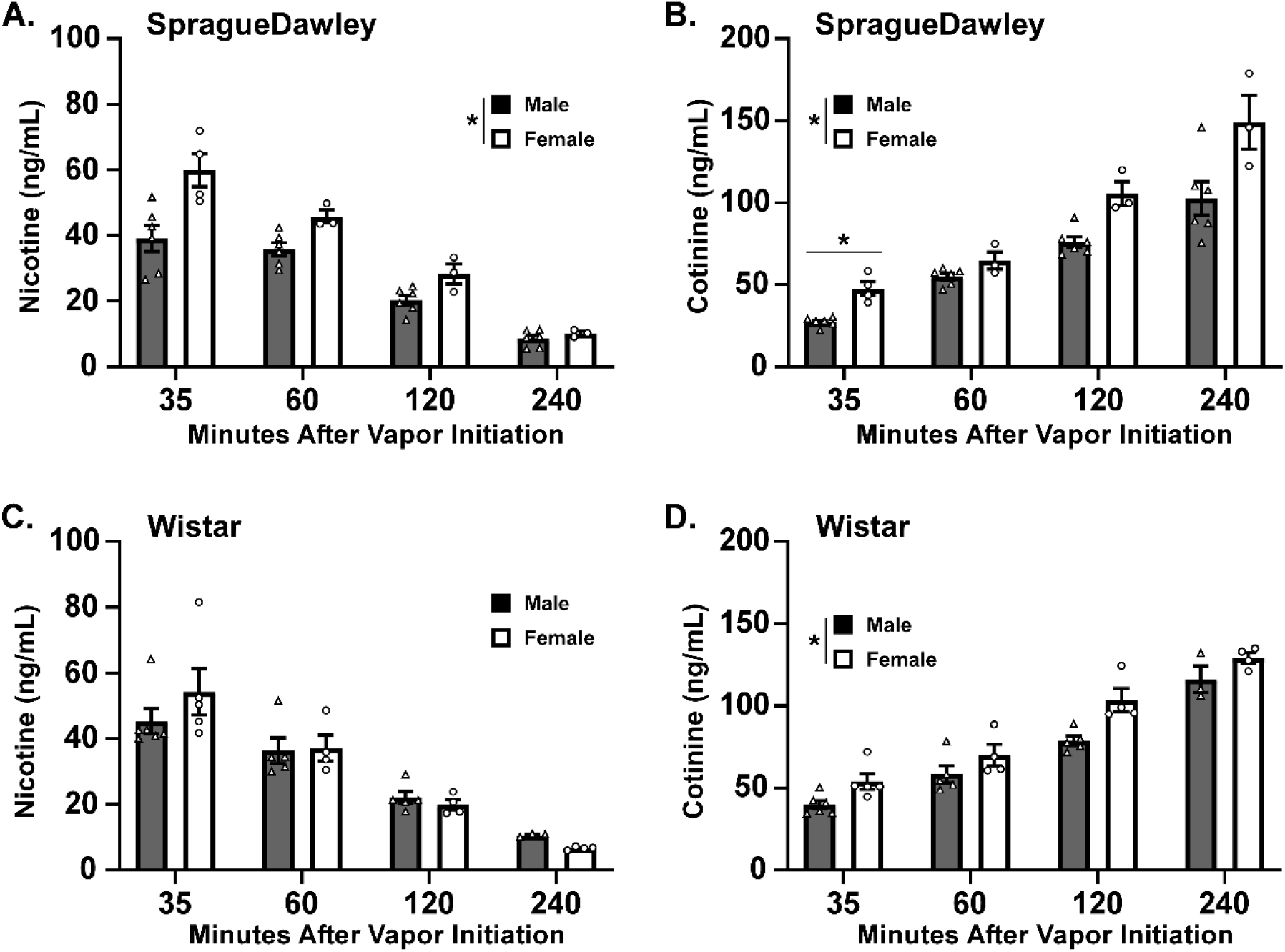
Mean (±SEM) and individual plasma concentrations of nicotine (A., C.) and cotinine (B., D.) for Male (N=6) and Female (N=4 35 min; N=3 60-240 min) Sprague-Dawley (A., B.) and Male (N=6 35 min; N=5 60-120; N=3 240) and Female (N=5 35 min; N=4 60-240 min) Wistar (C., D.) rats after vapor exposure to nicotine for 30 minutes. A significant difference between sexes, within strain, is indicated with *.

Follow-up analysis of cotinine was conducted across rat strain to determine any effect of sex [Sex: F (1, 19) = 24.46, P<0.0001; Time: F (1.259, 19.31) = 218.2, P<0.0001; Interaction: F (3, 46) = 3.472, P<0.05] and across sex to determine effects of strain [Strain: n.s.; Time: F (1.307, 20.04) = 178.9, P<0.0001; Interaction: n.s.]. Post-hoc analysis confirmed higher cotinine concentrations in the females, across strain, 35, 120 and 240 minutes after vapor initiation. Higher cotinine in female versus male Sprague-Dawley rats were also confirmed 35 minutes after vapor initiation. Post-hoc analysis further confirmed that plasma cotinine at each time-point significantly differed from the concentration at every other timepoint for male and female groups. A similar magnitude of sex difference in nicotine was observed after injection, however males exhibited the higher levels and the result did not reach statistical significance (**Supplemental Materials Figure S1**).

### Experiment 2: Effect of repeated adolescent nicotine inhalation on open field locomotion

#### Growth

There were no significant differences in body weight associated with the adolescent treatment (**Supplemental Materials Figure S3**).

#### Baseline locomotion

The initial three factor ANOVA confirmed that baseline locomotion in the open field was affected significantly by Sex [F (1, 28) = 8.67; P=0.01], by Time [F (11, 308) = 102.3; P<0.0001] and by the interaction of Time with Sex [F (11, 308) = 3.64; P<0.0001] (**Figure 2A**), but not by adolescent treatment (**Figure 2B**) nor any interaction of adolescent treatment with other factors. A follow up two-way ANOVA collapsed across adolescent treatment confirmed significant effects of Sex [F (1, 30) = 9.21; P<0.005], of Time [F (11, 330) = 102.6; P<0.0001] and of the interaction [F (11, 330) = 3.65; P<0.0001] on locomotor activity. The post-hoc test confirmed a sex difference in the 5-20 minute bins, and reduced activity compared with the first time bin was confirmed for all subsequent bins for all rats. Within the female group, activity was significantly lower in the 30-60-minute bins compared with the 15 minute timepoint and lower in the 30, 40-60 minute bins compared with the 20 minute timepoint. In the male group, however, activity was only significantly lower than the 15-minute observation in the final time bin. In the follow-up ANOVA collapsed across sex there was a significant effect of Time [F (11, 330) = 94.61; P<0.0001] on activity but not of adolescent treatment or the interaction of factors. The post-hoc test confirmed activity was significantly lower compared with the first time bin for all subsequent bins for both adolescent treatment groups.

**Figure 2:**
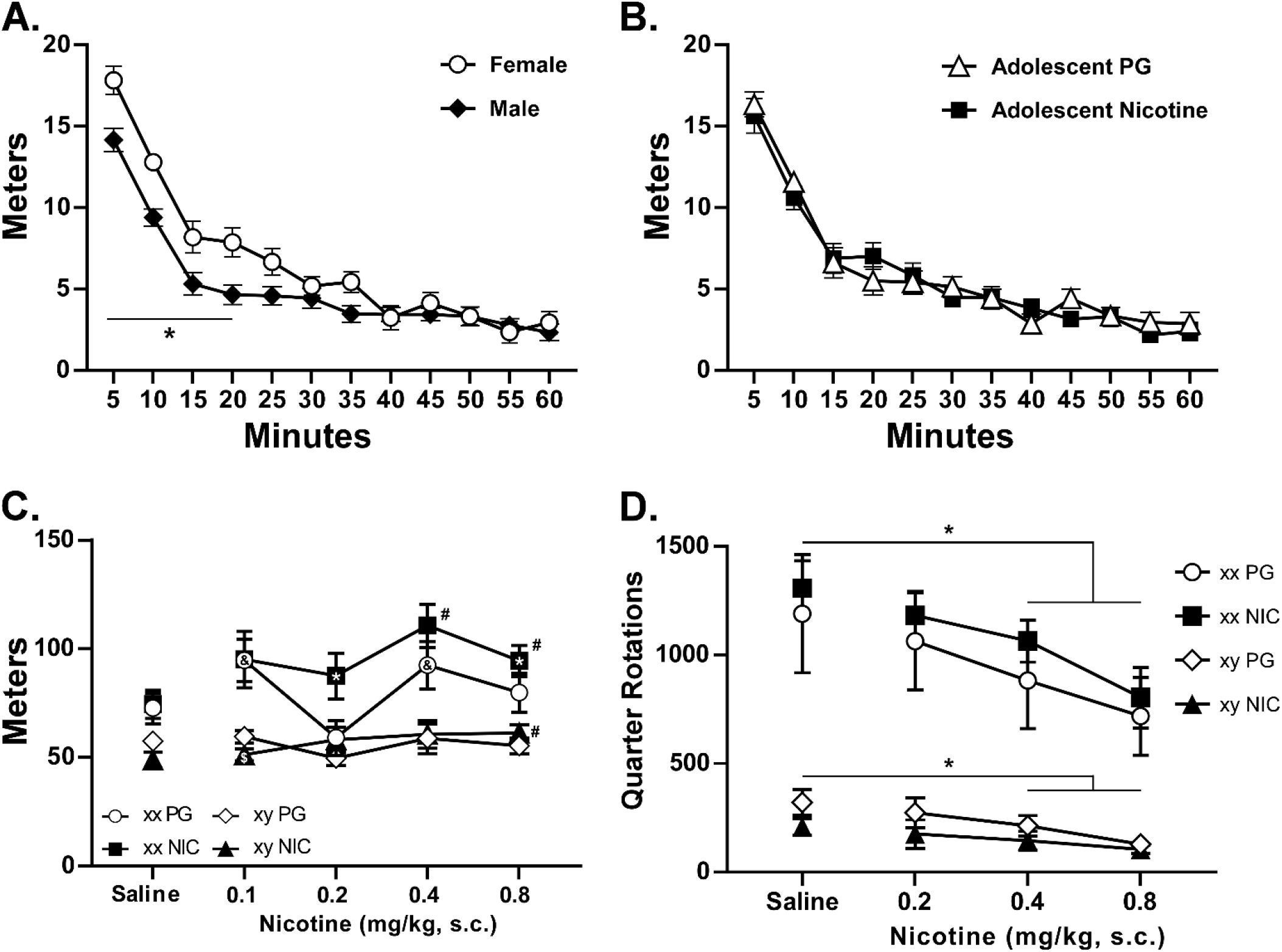
Mean (N=16 per group; ±SEM) distance traveled in the open arena by 5 minute time bin segments. The data are presented collapsed across A) adolescent treatment, and B) sex, to reflect the statistical analysis (see text). A significant difference between groups is indicated with *. C) Mean (N=8 per group; ±SEM) distance traveled in the open field after injection with saline or nicotine for male (xy) and female (xx) rats treated in adolescence with inhalation of vapor from the PG vehicle or NICotine (30 mg/mL). A significant difference from saline is indicated with #, from the 0.2 dose with &, from the 0.4 dose with * and from the 0.8 dose with $. D) Mean (±SEM) wheel activity after subcutaneous injection with saline or nicotine for male (N=8 per group) and female (N=8 per group) rats treated in adolescence with repeated inhalation of vapor from the propylene glycol (PG) vehicle or NICotine (30 mg/mL in the PG). A significant difference from Saline, within sex and collapsed across adolescent treatment, is indicated with *.

#### Nicotine injection

Open field distance was altered by nicotine (**Figure 2C**). The ANOVA confirmed significant effects of Dose [F (4, 112) = 11.39; P<0.0001], Sex [F (1, 28) = 23.62; P<0.0001] and the interactions of Dose with Sex [F (4, 112) = 5.51; P<0.0005] and Dose with Adolescent Treatment [F (4, 112) = 4.77; P<0.005] on distance traveled. The post-hoc test confirmed increased activity compared with saline injection in the female/Nicotine (0.4, 0.8 mg/kg) and male/Nicotine (0.8 mg/kg) groups. Activity was also higher in the female/Nicotine group after 0.4 mg/kg injection compared with either of the 0.2 or 0.8 mg/kg doses and higher in the male/Nicotine group after 0.8 mg/kg compared with the 0.1 mg/kg dose (**Figure 2C**). In the female/PG group, activity was significantly *reduced* after the 0.2 mg/kg dose compared with either of the 0.1 or 0.4 mg/kg doses. The effect of nicotine on activity by 5-minute bin is depicted in **Supplementary Materials Figure S4**.

### Experiment 3: Effect of repeated adolescent nicotine inhalation on wheel activity

Female rats engaged in more wheel activity than males (**Figure 2D**); see **Supplementary Materials Figure S5 for** baseline activity. Nicotine suppressed activity in both male and female rats, and to a similar extent across the adolescent exposure groups. The three-way ANOVA confirmed that total session wheel activity was significantly affected by Dose [F (2.682, 75.10) = 21.03; P<0.0001], by Sex [F (1, 28) = 46.10; P<0.0001] and by the interaction of Dose with Sex [F (3, 84) = 6.0; P<0.001]. There was no significant effect of Adolescent Treatment, nor of the interaction with other factors. Collapsed across adolescent treatment, there were significant effects of Dose [F (3, 90) = 22.17; P<0.0001], of Sex [F (1, 30) = 48.15; P<0.0001] and of the interaction [F (3, 90) = 6.32; P<0.001] confirmed. The post hoc test of the marginal mean for Dose confirmed activity was significantly lower after 0.8 mg/kg compared with all other conditions, and lower after 0.4 mg/kg compared with saline. This pattern was also confirmed within the female group, but no specific differences within the male group were confirmed in this analysis. Follow-up analysis of the male groups’ activity confirmed a significant effect of Dose [F (3, 42) = 9.64; P<0.0001], but not of Adolescent Treatment, and the post hoc test confirmed significantly lower wheel activity after 0.8 mg/kg compared with saline and the 0.2 mg/kg condition, as well as after 0.4 mg/kg compared with saline injection. Nicotine-induced reductions in wheel activity were most notable in the first 15-minutes (**Supplementary Materials Figure S6**).

### Experiment 4: Effect of repeated adolescent nicotine inhalation on nicotine vapor self-administration

#### Acquisition

Female rats obtained more nicotine (30 mg/mL) puffs than males in the first nine acquisition sessions and the tenth session (**Figure 3A, B**) conducted after four concentration-substitution sessions (**Figure 3C**). Vape deliveries during acquisition were significantly altered by rat Sex [F (1, 28) = 30.11; P<0.0001] and by the Session [F (9, 252) = 3.18; P<0.005] (**Figure 3A**). There were no significant effects of Adolescent treatment alone or in interaction with other factors confirmed. Collapsed across all groups, fewer deliveries were obtained in sessions 4, 7-10 relative to the first session as confirmed by the Dunnett post-hoc test. Each of the female groups obtained significantly more vapor deliveries than each of the male groups across sessions, but the treatment groups within each sex did not differ significantly from each other. The percentage of drug-associated responses was significantly altered by the Session [F (9, 252) = 3.98; P<0.0001] in the 3-way ANOVA (**Figure 3B**), but there were no significant effects of any other factor. Collapsed across group, rats expressed a higher percentage of drug-associated responses in sessions 5-10 relative to the first session, confirmed by the Dunnett post-hoc test.

**Figure 3:**
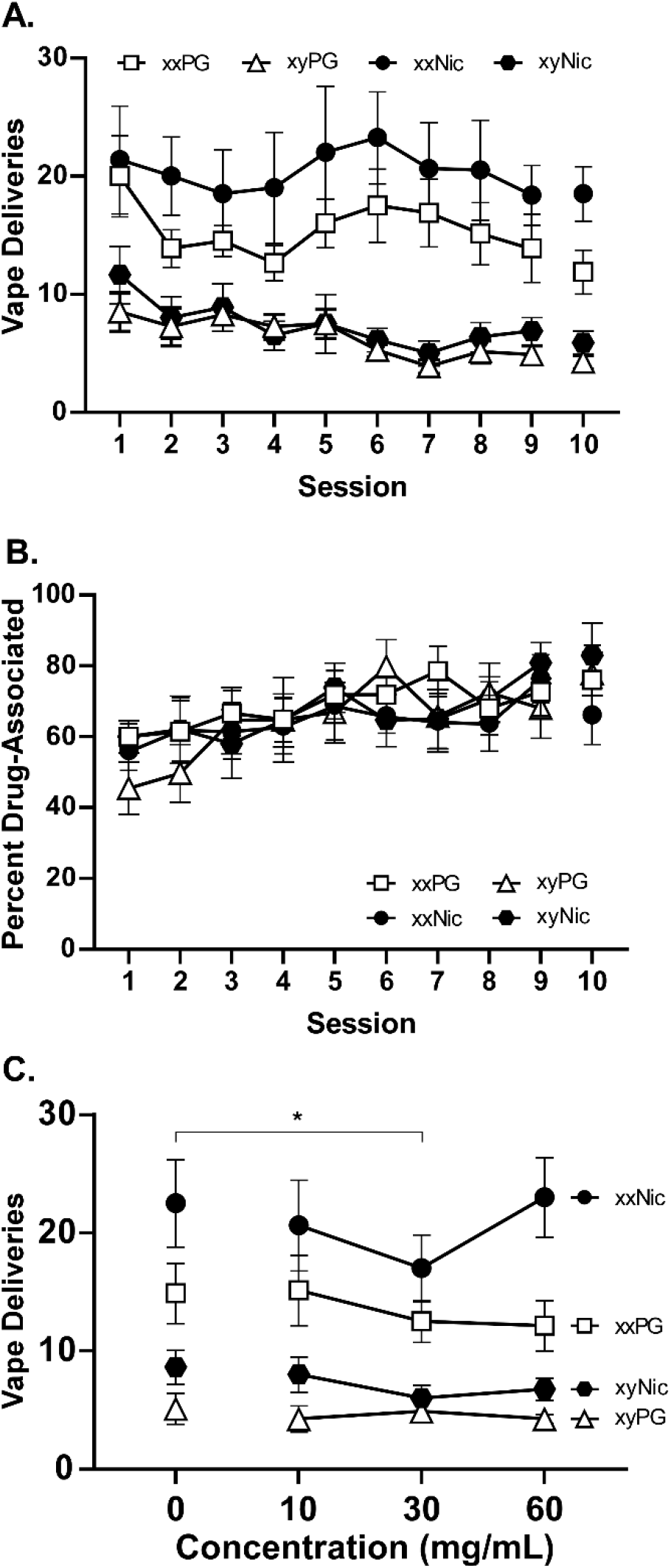
Mean (N=8 per group; ±SEM) A) nicotine (30 mg/mL) vapor deliveries, B) Percent responses on the drug-associated manipulandum in the initial nine 30-minute sessions and a 10^th^ session conducted after a 4-session dose-substitution. Intervals between sessions ranged from 2-5 days except for a 15-day interval between Session 4 and Session 5. C) Mean vapor deliveries obtained in the dose-substitution procedure. A significant difference from the 30 mg/mL training concentration, across groups, is indicated with *.

#### Concentration-Response

After acquisition session 9, rats were exposed to different concentrations of Nicotine (0, 10, 30, 60 mg/mL) in a counterbalanced order (**Figure 3C**). Female rats obtained more vapor deliveries than males, and the rats exposed to nicotine vapor as adolescents obtained more than the PG exposed groups. The three-factor ANOVA confirmed significant effects of Concentration [F (3, 84) = 2.74; P<0.05], Sex [F (1, 28) = 34.87; P<0.0001] and Adolescent Vapor exposure [F (1, 28) = 6.69; P<0.05] on vapor deliveries. The Tukey post-hoc analysis of the Concentration effect confirmed significant differences between the 30 mg/mL and 0 mg/mL concentrations. There were no significant differences in the percent of responses directed at the drug-associated manipulandum associated with any experimental factor (not shown). In the female groups, significantly more responses were made during the time-out on both the drug-associated and the alternate manipulandum when only PG vapor was available, compared with the nicotine 60 mg/mL vapor. More responses were also made on the alternate manipulandum when PG vapor was available, compared all three nicotine concentrations (**see Supplementary Materials Figure S7A, B**). The most nicotine-preferring female rats (by median split of the acquisition vapor deliveries) obtained more deliveries of 0 or 10 mg/mL vapor compared with the 30 mg/mL concentration. The lower half of the distribution obtained more deliveries of the 60 mg/mL nicotine vapor concentration compared with the 30 mg/mL concentration (**Supplementary Materials Figure S7C**).

### FR increment

<u>Deliveries of nicotine vapor</u> decreased compared with the immediately prior FR1 session (i.e., Session 10 in **Figure 3**) when the response contingency increased to FR5 (**Figure 4A**). The three-factor analysis confirmed that there was a significant effect of Session [F (3, 84) = 30.80; P<0.0001], of Sex [F (1, 28)v =29.54; P<0.0001], Adolescent treatment [F (1, 28) = 5.41; P<0.05], and of the interaction of Session with Sex [F (3, 84) = 6.66; P<0.0005] on vape deliveries. Post-hoc analysis further confirmed a difference between adolescent treatment conditions for the female rats on the FR1 and the third FR5 sessions. Vape deliveries were significantly lower in all of the FR5 sessions, compared with the FR1 session for both female groups and in the first FR5 session for the male nicotine-exposed group. Significantly *more* vape deliveries were obtained by the nicotine-exposed female group in the third FR5 session compared with the first FR5 session. <u>Responses on the drug-associated manipulandum</u> (**Figure 4B**) were significantly affected by Session [F (3, 84) = 23.28; P<0.0001], Sex [F (1, 28) = 23.34; P<0.0001], Adolescent treatment [F (1, 28) = 4.54; P<0.05], and the interaction of Session with Sex [F (3, 84) = 5.88; P<0.005]. The post-hoc test further confirmed that more responses on the drug-associated manipulandum were made by the PG-treated females (FR5 sessions 2 and 3) and the nicotine-treated females (all three FR5 sessions) compared with the FR1 session. Responses differed significantly between the two female groups in the third FR5 session, and the nicotine-exposed group emitted significantly more responses in the third FR5 session compared with the first FR5 session. There were no differences in <u>percent of responses directed at the drug-associated manipulandum</u> confirmed (**Figure 4C**). See **Supplementary Materials Figure S8** for the impact of restoring FR1 conditions after the interruption.

**Figure 4:**
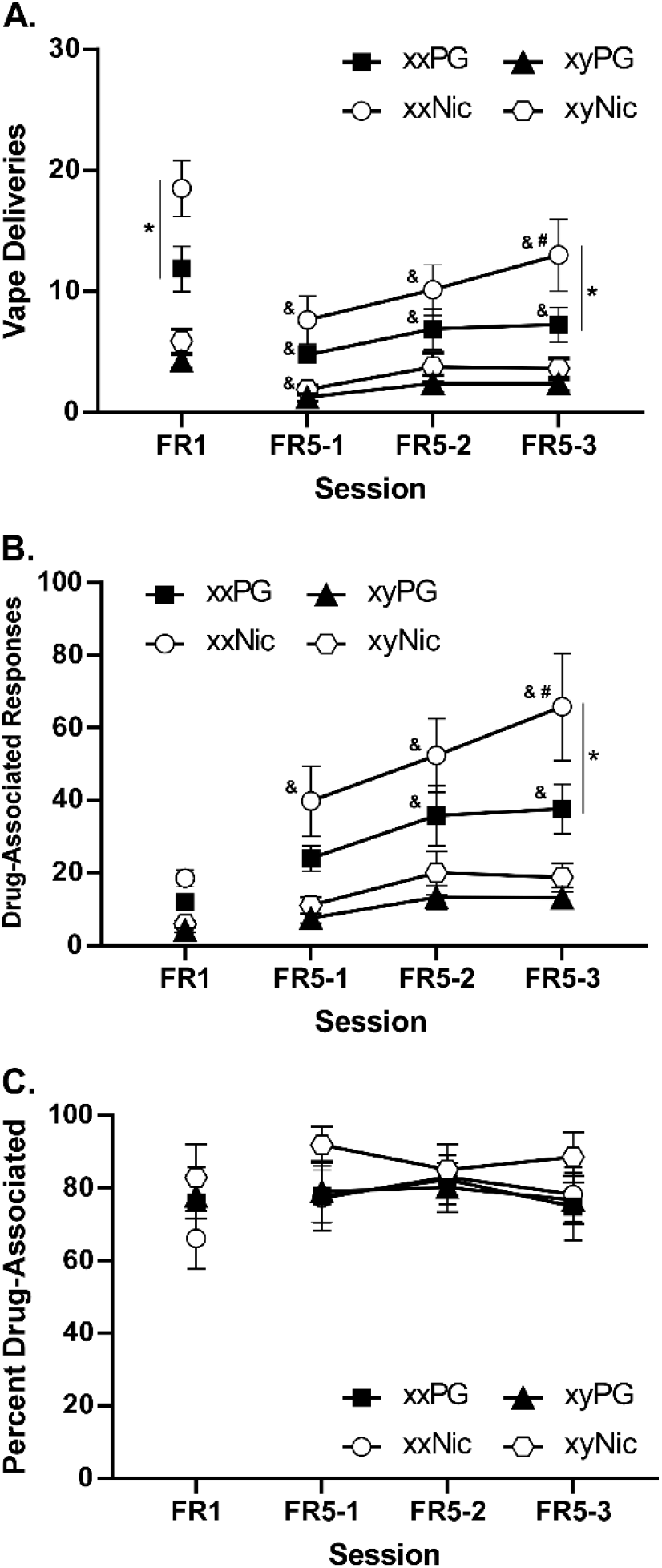
Mean (N=8 per group; ±SEM) A) nicotine (30 mg/mL) vapor deliveries, B) responses on the drug-associated manipulandum and C) percent responses on the drug-associated manipulandum obtained under Fixed Ratio (FR) 1 and then under FR 5 conditions for three sessions for 40 week old adult male (xy) and female (xx) rats exposed repeatedly during adolescence to vapor from the PG vehicle or <u>Nic</u>otine (30 mg/mL). A significant difference between treatment groups for a given session is indicated with *, a significant difference from the FR1 session within group with &, and a significant difference from the first FR5 session within group by #.

## Discussion

This study found that repeated adolescent nicotine vapor inhalation increases the locomotor stimulant effects of nicotine, and increases the vapor self-administration of nicotine, in female rats when they are evaluated as adults. No impact of this treatment on baseline locomotor activity in the open field or on wheels was observed. The study also showed that Electronic Nicotine Delivery System (ENDS) inhalation effectively delivers nicotine to male and female rats of two strains. There were only minor strain contributions to the concentrations of either cotinine or nicotine, but some evidence that female rats reach higher plasma levels for the same vaping conditions. The difference in effective dose may be relevant to any apparent sex differences observed. Furthermore, the temperature response of adolescent rats to nicotine vapor (**Supplementary Figure S2**) was similar to prior reports for adult rats. Thus, ENDS delivery of nicotine to rats generalizes well across sex, strain and age.

Females were consistently more active than males across telemetry (**Supplementary Figure S2**), open field and wheel assays. This is congruent with prior reports of activity measured with open field video or beam-break technology, radio-telemetry and activity wheels (Krentzel et al., 2020; Lee et al., 2019; Verlangieri, 1979). Nicotine injection significantly increased open field activity in the male (0.8 mg/kg, s.c.) and female (0.4, 0.8 mg/kg, s.c.) nicotine-exposed groups, but not the PG groups. Female and male rats were equivalently sensitive to the wheel activity suppressing effects of nicotine, expressing similar suppression in the first 15 min after 0.2-0.8 mg/kg, s.c., doses. Prior work found ∼0.25 mg/kg, s.c. to be the peak of an inverted-U dose response for increased wheel activity in female Sprague-Dawley rats (Gillman, Kosobud and Timberlake, 2010), showing that nicotine can either increase or decrease rodent activity depending on dose, assay and other factors.

Vapor *self-administration* procedures are relatively novel, although recent evidence demonstrates self-administration of nicotine (Lallai et al., 2021; Smith et al., 2020), heroin (Gutierrez et al., 2020), fentanyl (McConnell et al., 2021), sufentanil (Vendruscolo et al., 2018) and cannabis extracts (Freels et al., 2020). The present results show that male and female rats self-administer nicotine by vapor inhalation without any prior food-reinforced training (as in Lallai et al. 2021). Within group, rats obtained similar numbers of vapor deliveries under a FR1 contingency across the entire study. A higher response contingency (FR5) led to increased numbers of responses on the drug-associated manipulandum, which was associated with mean behavioral discrimination of over 75%, well over a 2:1 ratio used as evidence for self-administration in some studies (Gellner, Belluzzi and Leslie, 2016; Stringfield et al., 2023). A purely locomotor stimulant effect of nicotine (e.g. as in **Figure 2C** for the nicotine-exposed female rats) that also generically increased operant responses would increase interactions with the two manipulanda in equivalent number and therefore *reduce* behavioral discrimination. Female rats increased responding on both drug-associated and alternate manipulanda during the time-out interval when vehicle vapor was substituted and decreased responding on the drug-associated manipulandum when the higher concentration was delivered. Furthermore, a median split analysis shows that the more preferring half of the female distribution obtained more vapor in both 0 and 10 mg/mL substitution, whereas the lower preferring half of the distribution obtained more in the 60 mg/mL substitution, relative to the training concentration (**Supplementary Figure S7C**). Thus, sensitivity to nicotine concentration was exhibited albeit with individual differences in expression. Notably, the impact in the lower-preferring half would suggest that a higher training concentration may have produced higher mean self-administration in that subgroup. Overall, this pattern is consistent with drug-seeking behavior, i.e., self-administration.

There are caveats to this interpretation. The concentration-substitution did not produce strong differences in behavior across groups, and the main consistent difference was between the training concentration and increased responding for vehicle substitution. This latter is consistent with extinction bursting (Katz and Lattal, 2021), which in self-administration is a drug-seeking phenomenon produced by unexpected vehicle substitution, but could also be interpreted as responding for the vehicle. As shown in the Supplemental Materials, responding for PG vapor is initially high in naïve rats but undergoes extinction under a gradually increasing FR1 to FR5 contingency. Female rats reach stable session vapor delivery rates about half to a quarter of the rates observed in the current cohorts of female rats. In all there is strong, but not overwhelming, evidence for nicotine self-administration by vapor inhalation.

Males and females exhibited similar levels of behavioral discrimination, but females consistently obtained more vapor deliveries. Since there is considerable evidence that female rats will intravenously self-administer many drugs at a higher rate (Anker and Carroll, 2011), it is possible to view this outcome as a positive control that is consistent with drug self-administration. With respect to intravenous nicotine self-administration, a meta-analytic review of twenty studies concluded female rats self-administer nicotine at higher rates, with an estimate of the magnitude of effect of 0.18 standard deviations (Flores et al., 2019). There have, however, been a diversity of outcomes, which may interact with strain and other methodological variables, as is further described in the **Supplemental Discussion**. The impact of adolescent nicotine vapor exposure on self-administration was limited to the initial stages of the study for female rats. A somewhat variable group mean difference during acquisition led to stabilization of a significant difference in the initial FR5 experiment, since nicotine-exposed females self-administered more vapor deliveries in the baseline FR1 session and recovered more quickly from the introduction of FR5, leading to a mean difference on the third session.

Overall, these data do not illustrate significant lasting consequences of repeated adolescent nicotine exposure by vapor inhalation on baseline activity patterns in open field or on an exercise wheel. These data do, however, point to increased sensitivity to the locomotor stimulant effects of nicotine and a liability for moderately enhanced nicotine self-administration in adulthood, in female rats.

## Acknowledgements

The authors are grateful to Eric L. Harvey, Ph.D., for running plasma analysis studies.

## Literature Cited

Anker, J.J., Carroll, M.E., 2011. Females are more vulnerable to drug abuse than males: evidence from preclinical studies and the role of ovarian hormones. Curr Top Behav Neurosci 8, 73–96, doi: 10.1007/7854_2010_93.

Breit, K.R., Rodriguez, C.G., Hussain, S., Thomas, K.J., Zeigler, M., Gerasimidis, I., Thomas, J.D., 2022. A Model of Combined Exposure to Nicotine and Tetrahydrocannabinol via Electronic Cigarettes in Pregnant Rats. Front Neurosci 16, 866722, doi: 10.3389/fnins.2022.866722.

Cardenas, A., Lotfipour, S., 2022. Age- and Sex-Dependent Nicotine Pretreatment Effects on the Enhancement of Methamphetamine Self-administration in Sprague-Dawley Rats. Nicotine & tobacco research : official journal of the Society for Research on Nicotine and Tobacco 24(8), 1186–1192, doi: 10.1093/ntr/ntab218.

Chaudhri, N., Caggiula, A.R., Donny, E.C., Booth, S., Gharib, M.A., Craven, L.A., Allen, S.S., Sved, A.F., Perkins, K.A., 2005. Sex differences in the contribution of nicotine and nonpharmacological stimuli to nicotine self-administration in rats. Psychopharmacology 180(2), 258–266, doi: 10.1007/s00213-005-2152-3.

Chellian, R., Behnood-Rod, A., Wilson, R., Febo, M., Bruijnzeel, A.W., 2021. Adolescent nicotine treatment causes robust locomotor sensitization during adolescence but impedes the spontaneous acquisition of nicotine intake in adult female Wistar rats. Pharmacology, biochemistry, and behavior 207, 173224, doi: 10.1016/j.pbb.2021.173224.

Chellian, R., Behnood-Rod, A., Wilson, R., Kamble, S.H., Sharma, A., McCurdy, C.R., Bruijnzeel, A.W., 2020. Adolescent nicotine and tobacco smoke exposure enhances nicotine self-administration in female rats. Neuropharmacology 176, 108243, doi: 10.1016/j.neuropharm.2020.108243.

Cook, C.E., Jeffcoat, A.R., Hill, J.M., Pugh, D.E., Patetta, P.K., Sadler, B.M., White, W.R., Perez-Reyes, M., 1993. Pharmacokinetics of methamphetamine self-administered to human subjects by smoking S-(+)-methamphetamine hydrochloride. Drug Metab Dispos 21(4), 717–723, doi:

Cooper, S.Y., Akers, A.T., Henderson, B.J., 2021. Flavors Enhance Nicotine Vapor Self-administration in Male Mice. Nicotine & tobacco research : official journal of the Society for Research on Nicotine and Tobacco 23(3), 566–572, doi: 10.1093/ntr/ntaa165.

Diehl, K.H., Hull, R., Morton, D., Pfister, R., Rabemampianina, Y., Smith, D., Vidal, J.M., van de Vorstenbosch, C., 2001. A good practice guide to the administration of substances and removal of blood, including routes and volumes. J Appl Toxicol 21(1), 15–23, doi:

Echeveste Sanchez, M., Quadir, S.G., Whindleton, C.M., Hoffman, J.L., Faccidomo, S.P., Guhr Lee, T.N., Esther, C.R., Jr., Hodge, C.W., Herman, M.A., 2022. The effects of electronic nicotine vapor on voluntary alcohol consumption in female and male C57BL/6 J mice. Drug and alcohol dependence 241, 109676, doi: 10.1016/j.drugalcdep.2022.109676.

Espinoza, V.E., Giner, P., Liano, I., Mendez, I.A., O’Dell, L.E., 2022. Sex and age differences in approach behavior toward a port that delivers nicotine vapor. J Exp Anal Behav 117(3), 532–542, doi: 10.1002/jeab.756.

Flores, R.J., Uribe, K.P., Swalve, N., O’Dell, L.E., 2019. Sex differences in nicotine intravenous self-administration: A meta-analytic review. Physiology & behavior 203, 42–50, doi: 10.1016/j.physbeh.2017.11.017.

Fowler, C.D., Gipson, C.D., Kleykamp, B.A., Rupprecht, L.E., Harrell, P.T., Rees, V.W., Gould, T.J., Oliver, J., Bagdas, D., Damaj, M.I., Schmidt, H.D., Duncan, A., De Biasi, M., Basic Science Network of the Society for Research on, N., Tobacco, 2018. Basic Science and Public Policy: Informed Regulation for Nicotine and Tobacco Products. Nicotine & tobacco research : official journal of the Society for Research on Nicotine and Tobacco 20(7), 789–799, doi: 10.1093/ntr/ntx175.

Freels, T.G., Baxter-Potter, L.N., Lugo, J.M., Glodosky, N.C., Wright, H.R., Baglot, S.L., Petrie, G.N., Yu, Z., Clowers, B.H., Cuttler, C., Fuchs, R.A., Hill, M.N., McLaughlin, R.J., 2020. Vaporized Cannabis Extracts Have Reinforcing Properties and Support Conditioned Drug-Seeking Behavior in Rats. J Neurosci 40(9), 1897–1908, doi: 10.1523/JNEUROSCI.2416-19.2020.

Garber, J.C., Barbee, R.W., Bielitzki, J.T., Clayton, L.A., Donovan, J.C., Hendriksen, C.F.M., Kohn, D.F., Lipman, N.S., Locke, P.A., Melcher, J., Quimby, F.W., Turner, P.V., Wood, G.A., Wurbel, H., 2011. Guide for the Care and Use of Laboratory Animals, 8th Edition. National Academies Press, Washington D.C.

Gellner, C.A., Belluzzi, J.D., Leslie, F.M., 2016. Self-administration of nicotine and cigarette smoke extract in adolescent and adult rats. Neuropharmacology 109, 247–253, doi: 10.1016/j.neuropharm.2016.06.026.

Gillman, A.G., Kosobud, A.E., Timberlake, W., 2010. Effects of multiple daily nicotine administrations on pre- and post-nicotine circadian activity episodes in rats. Behav Neurosci 124(4), 520–531, doi: 10.1037/a0020272.

Gilpin, N.W., Wright, M.J., Jr., Dickinson, G., Vandewater, S.A., Price, J.U., Taffe, M.A., 2011. Influences of activity wheel access on the body temperature response to MDMA and methamphetamine. Pharmacology, biochemistry, and behavior 99(3), 295–300, doi: 10.1016/j.pbb.2011.05.006.

Gutierrez, A., Creehan, K.M., Taffe, M.A., 2021. A vapor exposure method for delivering heroin alters nociception, body temperature and spontaneous activity in female and male rats. J Neurosci Methods 348, 108993, doi: 10.1016/j.jneumeth.2020.108993.

Gutierrez, A., Nguyen, J.D., Creehan, K.M., Taffe, M.A., 2020. Female rats self-administer heroin by vapor inhalation. Pharmacology, biochemistry, and behavior 199, 173061, doi: 10.1016/j.pbb.2020.173061.

Henderson, B.J., Cooper, S.Y., 2021. Nicotine formulations impact reinforcement-related behaviors in a mouse model of vapor self-administration. Drug and alcohol dependence 224, 108732, doi: 10.1016/j.drugalcdep.2021.108732.

Hussain, S., Breit, K.R., Thomas, J.D., 2022. The effects of prenatal nicotine and THC E-cigarette exposure on motor development in rats. Psychopharmacology 239(5), 1579–1591, doi: 10.1007/s00213-022-06095-8.

Javadi-Paydar, M., Creehan, K.M., Kerr, T.M., Taffe, M.A., 2019a. Vapor inhalation of cannabidiol (CBD) in rats. Pharmacology, biochemistry, and behavior 184, 172741, doi: 10.1016/j.pbb.2019.172741.

Javadi-Paydar, M., Kerr, T.M., Harvey, E.L., Cole, M., Taffe, M.A., 2019b. Effects of nicotine and THC vapor inhalation administered by an electronic nicotine delivery system (ENDS) in male rats. Drug and alcohol dependence 198, 54–62, doi: 10.1016/j.drugalcdep.2019.01.027.

Johnston, L.D., Miech, R.A., O’Malley, P.M., Bachman, J.G., Schulenberg, J.E., Patrick, M.E., 2021. Monitoring the Future national survey results on drug use, 1975-2020: 2020 Overview Key Findings on Adolescent Drug Use. http://www.monitoringthefuture.org/pubs/monographs/mtf-overview2020.pdf.

Katz, B.R., Lattal, K.A., 2021. What is an extinction burst?: A case study in the analysis of transitional behavior. J Exp Anal Behav 115(1), 129–140, doi: 10.1002/jeab.642.

Khalouzadeh, F., Azizi, H., Semnanian, S., 2022. Adolescent nicotine exposure increases nociceptive behaviors in rat model of formalin test: Involvement of ventrolateral periaqueductal gray neurons. Life Sci 299, 120551, doi: 10.1016/j.lfs.2022.120551.

Krentzel, A.A., Proano, S., Patisaul, H.B., Meitzen, J., 2020. Temporal and bidirectional influences of estradiol on voluntary wheel running in adult female and male rats. Horm Behav 120, 104694, doi: 10.1016/j.yhbeh.2020.104694.

Lallai, V., Chen, Y.C., Roybal, M.M., Kotha, E.R., Fowler, J.P., Staben, A., Cortez, A., Fowler, C.D., 2021. Nicotine e-cigarette vapor inhalation and self-administration in a rodent model: Sex- and nicotine delivery-specific effects on metabolism and behavior. Addict Biol 26(6), e13024, doi: 10.1111/adb.13024.

Lee, J.R., Tapia, M.A., Nelson, J.R., Moore, J.M., Gereau, G.B., Childs, T.E., Vieira-Potter, V.J., Booth, F.W., Will, M.J., 2019. Sex dependent effects of physical activity on diet preference in rats selectively bred for high or low levels of voluntary wheel running. Behavioural brain research 359, 95–103, doi: 10.1016/j.bbr.2018.10.018.

Levin, E.D., Slade, S., Wells, C., Cauley, M., Petro, A., Vendittelli, A., Johnson, M., Williams, P., Horton, K., Rezvani, A.H., 2011. Threshold of adulthood for the onset of nicotine self-administration in male and female rats. Behavioural brain research 225(2), 473–481, doi: 10.1016/j.bbr.2011.08.005.

McConnell, S.A., Brandner, A.J., Blank, B.A., Kearns, D.N., Koob, G.F., Vendruscolo, L.F., Tunstall, B.J., 2021. Demand for fentanyl becomes inelastic following extended access to fentanyl vapor self-administration. Neuropharmacology 182, 108355, doi: 10.1016/j.neuropharm.2020.108355.

Miller, M.L., Moreno, A.Y., Aarde, S.M., Creehan, K.M., Vandewater, S.A., Vaillancourt, B.D., Wright, M.J., Jr., Janda, K.D., Taffe, M.A., 2013. A methamphetamine vaccine attenuates methamphetamine-induced disruptions in thermoregulation and activity in rats. Biol Psychiatry 73(8), 721–728, doi: 10.1016/j.biopsych.2012.09.010.

Montanari, C., Kelley, L.K., Kerr, T.M., Cole, M., Gilpin, N.W., 2020. Nicotine e-cigarette vapor inhalation effects on nicotine & cotinine plasma levels and somatic withdrawal signs in adult male Wistar rats. Psychopharmacology 237(3), 613–625, doi: 10.1007/s00213-019-05400-2.

Nguyen, J.D., Aarde, S.M., Cole, M., Vandewater, S.A., Grant, Y., Taffe, M.A., 2016. Locomotor Stimulant and Rewarding Effects of Inhaling Methamphetamine, MDPV, and Mephedrone via Electronic Cigarette-Type Technology. Neuropsychopharmacology : official publication of the American College of Neuropsychopharmacology 41(11), 2759–2771, doi: 10.1038/npp.2016.88.

Nguyen, J.D., Creehan, K.M., Kerr, T.M., Taffe, M.A., 2020. Lasting effects of repeated (9) - tetrahydrocannabinol vapour inhalation during adolescence in male and female rats. British journal of pharmacology 177(1), 188–203, doi: 10.1111/bph.14856.

Nguyen, J.D., Grant, Y., Creehan, K.M., Hwang, C.S., Vandewater, S.A., Janda, K.D., Cole, M., Taffe, M.A., 2019. Delta(9)-tetrahydrocannabinol attenuates oxycodone self-administration under extended access conditions. Neuropharmacology 151, 127–135, doi: 10.1016/j.neuropharm.2019.04.010.

Nguyen, J.D., Grant, Y., Taffe, M.A., 2021. Paradoxical changes in brain reward status during oxycodone self-administration in a novel test of the negative reinforcement hypothesis. British journal of pharmacology 178(18), 3797–3812, doi: 10.1111/bph.15520.

Nguyen, J.D., Hwang, C.S., Grant, Y., Janda, K.D., Taffe, M.A., 2018. Prophylactic vaccination protects against the development of oxycodone self-administration. Neuropharmacology 138, 292–303, doi: 10.1016/j.neuropharm.2018.06.026.

Park-Lee, E., Ren, C., Cooper, M., Cornelius, M., Jamal, A., Cullen, K.A., 2022. Tobacco Product Use Among Middle and High School Students - United States, 2022. MMWR Morb Mortal Wkly Rep 71(45), 1429–1435, doi: 10.15585/mmwr.mm7145a1.

Renda, B., Andrade, A.K., Frie, J.A., Sgarbossa, C.L., Murray, J.E., Khokhar, J.Y., 2020. High-dose adolescent nicotine exposure permits spontaneous nicotine self-administration in adult male rats. Drug and alcohol dependence 215, 108215, doi: 10.1016/j.drugalcdep.2020.108215.

Schulenberg, J.E., Patrick, M.E., Johnston, L.D., O’Malley, P.M., Bachman, J.G., Miech, R.A., 2021. Monitoring the Future national survey results on drug use, 1975-2020. Volume 2, College students and adults ages 19–60.

Smith, L.C., Kallupi, M., Tieu, L., Shankar, K., Jaquish, A., Barr, J., Su, Y., Velarde, N., Sedighim, S., Carrette, L.L.G., Klodnicki, M., Sun, X., de Guglielmo, G., George, O., 2020. Validation of a nicotine vapor self-administration model in rats with relevance to electronic cigarette use. Neuropsychopharmacology : official publication of the American College of Neuropsychopharmacology 45(11), 1909–1919, doi: 10.1038/s41386-020-0734-8.

Stringfield, S.J., Sanders, B.E., Suppo, J.A., Sved, A.F., Torregrossa, M.M., 2023. Nicotine Enhances Intravenous Self-administration of Cannabinoids in Adult Rats. Nicotine & tobacco research : official journal of the Society for Research on Nicotine and Tobacco 25(5), 1022–1029, doi: 10.1093/ntr/ntac267.

Taffe, M.A., Creehan, K.M., Vandewater, S.A., Kerr, T.M., Cole, M., 2021. Effects of Delta(9)-tetrahydrocannabinol (THC) vapor inhalation in Sprague-Dawley and Wistar rats. Experimental and clinical psychopharmacology 29(1), 1–13, doi: 10.1037/pha0000373.

Vendruscolo, J.C.M., Tunstall, B.J., Carmack, S.A., Schmeichel, B.E., Lowery-Gionta, E.G., Cole, M., George, O., Vandewater, S.A., Taffe, M.A., Koob, G.F., Vendruscolo, L.F., 2018. Compulsive-Like Sufentanil Vapor Self-Administration in Rats. Neuropsychopharmacology : official publication of the American College of Neuropsychopharmacology 43(4), 801–809, doi: 10.1038/npp.2017.172.

Verlangieri, A.J., 1979. Prenatal and postnatal chronic lead intoxication and running wheel activity in the rat. Pharmacology, biochemistry, and behavior 11(1), 95–98, doi: 10.1016/0091-3057(79)90303-4.

