## Supplementary Materials for "Adult consequences of repeated nicotine vapor inhalation in adolescent rats"

### Supplementary Methods

#### *Subjects*

Additional groups of male (N=8) and female (N=10) Wistar (CRL) rats were used for further investigation of operant responding for vehicle vapor. The vivarium was kept on a 12:12 hour reversed light-dark cycle, and behavior studies were conducted during the vivarium dark period. Food and water were provided ad libitum in the home cage. Procedures were conducted in accordance with protocols approved by the Institutional Animal Care and Use Committee of The Scripps Research Institute and were consistent with the NIH Guide (Garber et al., 2011).

**Catheterization:** Rats for the blood sampling study were anesthetized with an isoflurane/oxygen vapor mixture (isoflurane 5% induction, 1-3% maintenance) and prepared with chronic indwelling intravenous catheters. The catheters consisted of a 14.5-cm length of polyurethane based tubing (Micro-Renathane®, Braintree Scientific, Inc, Braintree, MA) fitted to a guide cannula (Plastics One, Roanoke, VA) curved at an angle and encased in dental cement anchored to an ~3 cm circle of durable mesh. Catheter tubing was passed subcutaneously from the animal's back to the right jugular vein. Catheter tubing was inserted into the vein and tied gently with suture thread. A liquid tissue adhesive was used to close the incisions (3M™ Vetbond™ Tissue Adhesive: 1469SB, 3M, St. Paul, MN). A minimum of 7 days was allowed for surgical recovery prior to starting an experiment. For the first three days of the recovery period, an antibiotic (cefazolin) and an analgesic (flunixin) were administered daily. Catheters were flushed with ~0.2-0.3 ml heparinized (166.7 USP/ml) saline before sessions and ~0.2-0.3 ml heparinized saline containing cefazolin (100 mg/mL) after sessions.

#### *Plasma Nicotine and Cotinine Analysis*

Plasma nicotine and cotinine concentrations were quantified using liquid chromatography/mass spectrometry (LCMS) as previously reported (Javadi-Paydar, Mehrak et al., 2019). Briefly, 50 ul of plasma were mixed with 50 ul of deuterated internal standard (100 ng/ml cotinine-d3 and nicotine-d4; Cerilliant). Nicotine and cotinine (and the internal standards) were extracted into 900 ul of acetonitrile and then dried. Samples were reconstituted in 100 uL of an acetonitrile/water (9:1) mixture. Separation was performed on an Agilent LC1200 with an Agilent Poroshell 120 HILIC column (2.1mm x 100mm; 2.7 um) using an isocratic mobile phase composition of acetonitrile/water (90:10) with 0.2% formic acid at a flow rate of 325 uL/min. Nicotine and cotinine were quantified using an Agilent MSD6130 single quadrupole interfaced with electrospray ionization and selected ion monitoring [nicotine (m/z=163.1), nicotine-d4 (m/z=167.1), cotinine (m/z=177.1) and cotinine-d3 (m/z=180.1)]. Calibration curves were generated daily using a concentration range of 0-200 ng/mL with observed correlation coefficients of 0.999.

#### *Apparatus*

Vapor Inhalation: An e-cigarette based vapor inhalation system (La Jolla Alcohol Research, Inc) which has previously been shown to deliver active doses of $\Delta^9$-tetrahydrocannabinol (THC), cannabidiol, heroin, oxycodone, methamphetamine and nicotine (Gutierrez, Creehan and Taffe, 2021; Javadi-Paydar, M. et al., 2019a; Javadi-Paydar, M. et al., 2019b; Nguyen et al., 2016; Nguyen et al., 2019) was used for these studies. Vapor was delivered into sealed vapor exposure chambers (209 mm W X 234 mm H X 259 mm L, passive vapor exposure; 152 mm W X 178 mm H X 330 mm L, vapor self-administration; La Jolla Alcohol Research, Inc, La Jolla, CA, USA) through the use of e-vape controllers (Model SSV-3 or SVS-200; 58 watts, 0.24-0.26 ohms, 3.95-4.3 volts, ~214 °F; La Jolla Alcohol Research, Inc, La Jolla, CA, USA) to trigger SMOK Baby Beast Brother TFV8 sub-ohm tanks. Tanks were equipped with V8 X-Baby M2 0.25 ohm coils. MedPC IV software was used to schedule and trigger vapor delivery (Med Associates, St. Albans, VT USA). For passive vapor exposure, 6 second vapor puffs were triggered at 5-minute intervals. The apparatus and settings were the same for all drug conditions. The chamber air was vacuum-controlled by a chamber exhaust valve (i.e., a "pull" system) to flow room ambient air through an intake valve at ~1 L per minute. This also functioned to ensure that vapor entered the chamber on each device triggering event. The vapor stream was integrated with the ambient air stream once triggered. Airflow was initiated 30 seconds prior to, and discontinued 10 seconds after, each puff. For vapor self-administration, air flowed continuously at ~2 L per minute, and 1 second vapor puffs were triggered for each reinforcer delivery.

Open Field: Testing was conducted in plastic chambers with dimensions of 80 cm (L) X 44 cm (W) X 33 cm (H). Locomotor activity was recorded using webcams (Logitech Model C270) mounted approximately 1 meter above the arena. Analysis of behavior was conducted off-line using ANY-maze behavior tracking software (Stoelting) with analysis configured for the Center (43 cm X 11 cm) versus the Peripheral Zones. The experiment was run under white light illumination to facilitate the video tracking.

Activity Wheels: Experimental sessions were conducted in white illuminated procedure rooms with activity wheels that attached to a typical housing chamber with a door cut to provide access to the wheel (Med Associates; Model ENV-046), using approaches previously described (Gilpin et al., 2011; Miller et al., 2013; Taffe et al., 2021). The lights were turned on for maximum comparison with the open field experiment. Rats were given access to the wheel in acute 30-minute sessions during which wheel rotation (quarter-rotation resolution) activity was recorded at 10-minute intervals. One 30-minute habituation session was conducted for each animal prior to initiating the experimental sessions.

*Nicotine inhalation in adolescent rats*

Studies were conducted in groups of male (N=8) and female (N=8) peri-adolescent Sprague-Dawley rats, separate from the main study adolescents, to verify efficacy of the nicotine vapor inhalation procedure in younger animals since our prior studies (Javadi-Paydar, M. et al., 2019a; Javadi-Paydar, Mehrak et al., 2019) had been conducted in adults. Rats arrived in the laboratory on Post-Natal Day (PND) 22 and were implanted with radio-telemetry transmitters (Data Sciences International, St. Paul, MN, USA; TA11TA-F20) on PND 31 using aseptic technique, inhalation anesthesia and post-operative recovery as previously described (Nguyen et al., 2020; Taffe, Creehan and Vandewater, 2015). One animal of each sex did not survive through surgical recovery, thus N=7 per group for the studies. Vapor experiments were initiated on PND 43 with a baseline assessment, the first set of nicotine vapor experiments were conducted on PND 49 and PND 52, THC vapor inhalation experiments not herein reported on PND 57 and PND 60, and then a repetition of nicotine on PND 63 and PND 66. The goal was to determine if body temperature responses to nicotine vapor in late adolescence are similar to those reported for the adult age range, using identical vapor inhalation conditions (Javadi-Paydar, Mehrak et al., 2019). In brief, rats were placed in individual chambers for baseline recording for 30 minutes. Thereafter they were placed in an inhalation chamber for the 30 minute exposure period, and returned to the individual telemetry recording chamber for post-inhalation evaluation. The nicotine experiments determined the effects of PG inhalation versus nicotine (30 mg/mL in the PG) inhalation, in a counterbalanced order within groups, on spontaneous activity and body temperature responses. Three males had a significantly aberrant drop in body temperature following nicotine vapor on the same experimental day for the first experiment, and it was noted that room temperature was ~0.2° C colder than for any other study day. For this reason the nicotine experiment was repeated a second time. Comparison of the temperature and activity means with those three animals excluded with the full sample results from the repetition study confirmed qualitative similarity of results for both male and female rats. Thus, these pre-planned inhalation conditions (nicotine 30 mg/mL for 30 minutes) were used for Experiment 2. Subsequent experiments conducted in this group of animals have been previously reported (Gutierrez, Creehan and Taffe, 2021).

*PG Vapor Self-Administration / Extinction*

Male and female Wistar rats were permitted to respond for PG vapor deliveries in one hour sessions, starting at PND 90-105. The male group was limited to 5 reinforcer deliveries for the first two sessions but otherwise no restrictions were imposed. The male group's response contingency was incremented to FR2 on Session 7, to FR3 on Session 12 and to FR5 on Session 16. The female group's response contingency was incremented to FR3 on Session 6 and to FR5 on Session 11.

*Data Analysis*

Telemetry data were analyzed by 5-minute bin with a within-subject factor of Time after the start of inhalation, and a factor for Group, included in the ANOVA. The telemetry analysis included three pre-inhalation baseline values and data from 40-120 minutes after the start of inhalation.

### Supplementary Results

*Experiment 1: Plasma nicotine and cotinine after subcutaneous injection*

The female rats with patent catheters were too few for the subcutaneous injection study, thus it was opted to collect samples 15 and 120 minutes after injection by acute venipuncture under inhalation anesthesia (with recovery in-between sampling points). Statistical comparisons were therefore limited to the 15- and 120-minute samples, however the intervening time points for the males are also presented in **Figure S1** for reference. The mixed-effects analysis of all four groups confirmed a significant effect of Time post-injection [F (1, 18) = 35.78, P<0.0001] but not of group for nicotine concentrations. This was also the outcome when the analysis was limited to each strain, to each sex, collapsed across strain or collapsed across sex. The mixed-effects analysis of all four groups confirmed a significant effect of Time post-injection [F (1, 18) = 35.78, P<0.0001] and of Group [F (3, 19) = 4.58; P<0.05] for cotinine concentrations. The post-hoc test confirmed that there was a significant difference in cotinine concentrations between strains within both male and female groups at the 120-minute time point. There was also a significant difference between Sprague-Dawley males and Wistar females at this-time point.

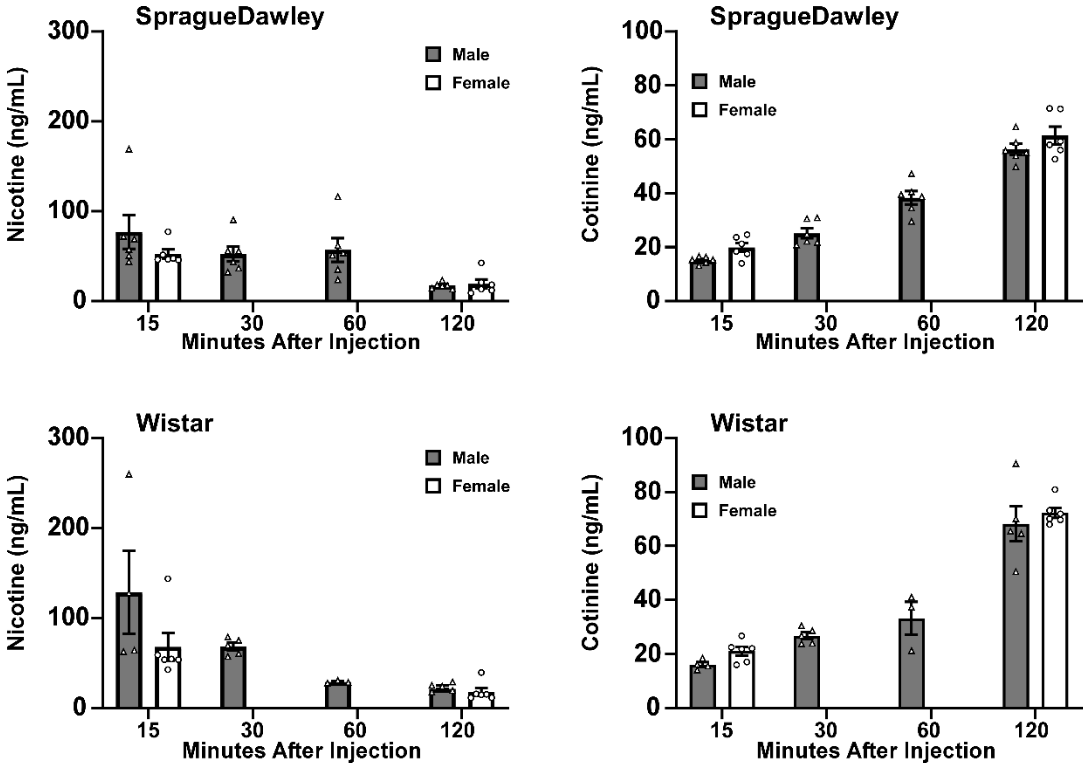


***Figure S1:*** *Mean (±SEM) and individual plasma nicotine and cotinine concentrations observed in male (N=6) and female (N=6) Sprague-Dawley and in male (N=4-5 per time point out of N=5 total rats) and female (N=6) Wistar rats, after 1 mg/kg, nicotine s.c..*

<u>*Nicotine inhalation in adolescent rats*</u>

There was a modest and short-lived reduction in body temperature observed in both sexes of peri-adolescent rats after nicotine inhalation (**Figure S2**), similar to what we reported previously for adult male Sprague-Dawley rats after a single 30 min nicotine vapor exposure (Javadi-Paydar, Mehrak et al., 2019), thus confirming active doses are delivered by this technique in adolescent animals. No significant effects on activity rate were observed. The two-factor analysis confirmed significant effects of Group [F (3, 24) = 9.31; P<0.0005] and the interaction of Group with Time after vapor initiation [F (57, 456) = 5.40; P<0.0001] on body temperature.

The post-hoc analysis further confirmed that body temperature was lower after nicotine inhalation compared with PG inhalation in female (40-55 minutes after the start of inhalation)

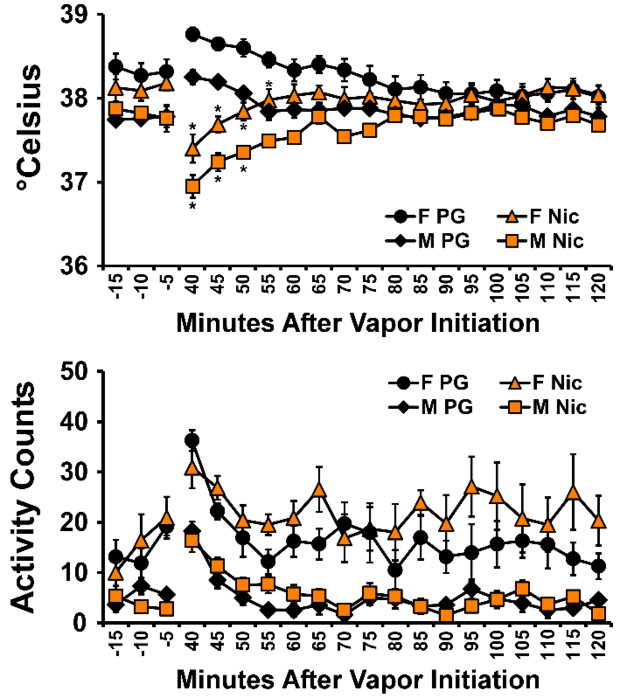


***Figure S2**: Mean (±SEM) temperature and activity rates for male (N=7) and female (N=7) rats assessed after inhalation of vapor from the propylene glycol (PG) vehicle or nicotine (30 mg/mL in the PG) on PND 63 and 66. A significant difference between inhalation conditions, within sex, is indicated with *.*

and male (40-50 minutes after the start of inhalation) rats. The post-hoc test likewise confirmed that male rat temperature was lower than female rat temperature after PG (all three baseline samples, 40, 50-65 minutes after vapor initiation) or nicotine (50-60 minutes after vapor initiation) inhalation.

The two-factor analysis of activity confirmed significant effects of Group [F (3, 24) = 22.42; P<0.0001] and of Time [F (19, 456) = 6.96; P<0.0001], but not the interaction of factors, on activity rates. The post-hoc test of the marginal means for Group confirmed that female activity was higher in each treatment condition compared with male activity in each treatment condition, however there was no significant difference in activity associated with treatment condition within either sex.

***Experiment 2: Effect of repeated adolescent nicotine inhalation on open field locomotion in adult rats***

Growth:

There was no immediate or lasting impact of the adolescent vapor exposure on bodyweight (**Figure S3**).

Locomotor Activity:

In the Open Field experiments, the female rats traveled more distance than did the males, particularly in the first five to ten minutes of the sessions (**Figure S4**). There were significant effects of Sex and session Time confirmed for the first Saline [Time: F (11, 308) = 129.5; P<0.0001; Sex: F (1, 28) = 17.62;

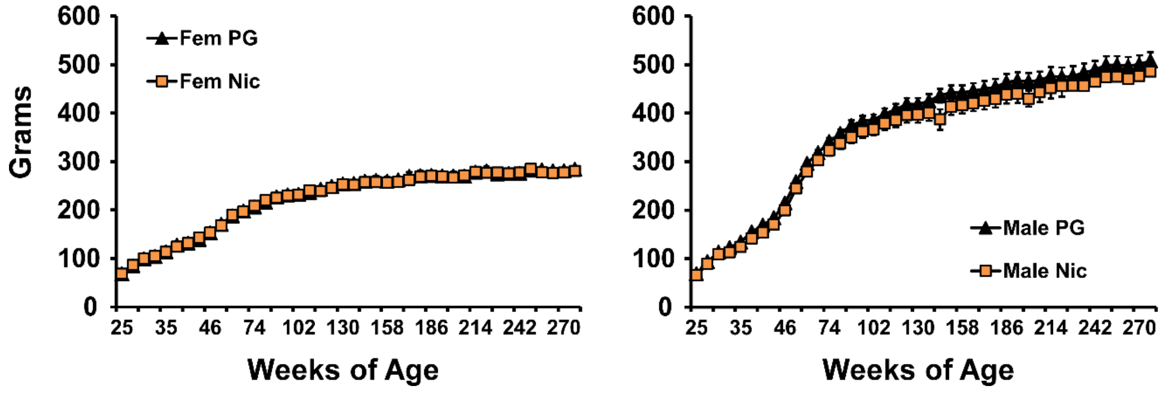


***Figure S3:** Mean (N=8 per group; ±SEM) body weight for Female and Male rats exposed to vapor from the PG vehicle, or Nicotine during adolescence.*

P<0.0005; Interaction of Time with Sex: F (11,308) = 2.18; P<0.05] and second Saline [Time: F (11, 297) = 130.9; P<0.0001; Sex: F (1, 27) = 10.63; P<0.005; Interaction of Time with Sex: F (11, 297) = 2.00; P<0.05] sessions. Consistent effects of adolescent treatment condition were also observed in the 0.1 mg/kg [Time: F (11, 308) = 152.7; P<0.0001; Sex: F (1, 28) = 22.50; P<0.0001; Interaction of Time with Sex: F (11, 308) = 2.18; P<0.05; Interaction of Time, Sex and Adolescent treatment: F (11, 308) = 1.90; P<0.05] and the 0.2 mg/kg [Time: F (11, 308) = 28.78; P<0.0001; Sex: F (1, 28) = 6.84; P<0.05; Adolescent treatment: F (1, 28) = 6.13; P<0.05; Interaction of Time with Sex: F (11, 308) = 2.02; P<0.05] nicotine treatment studies. No

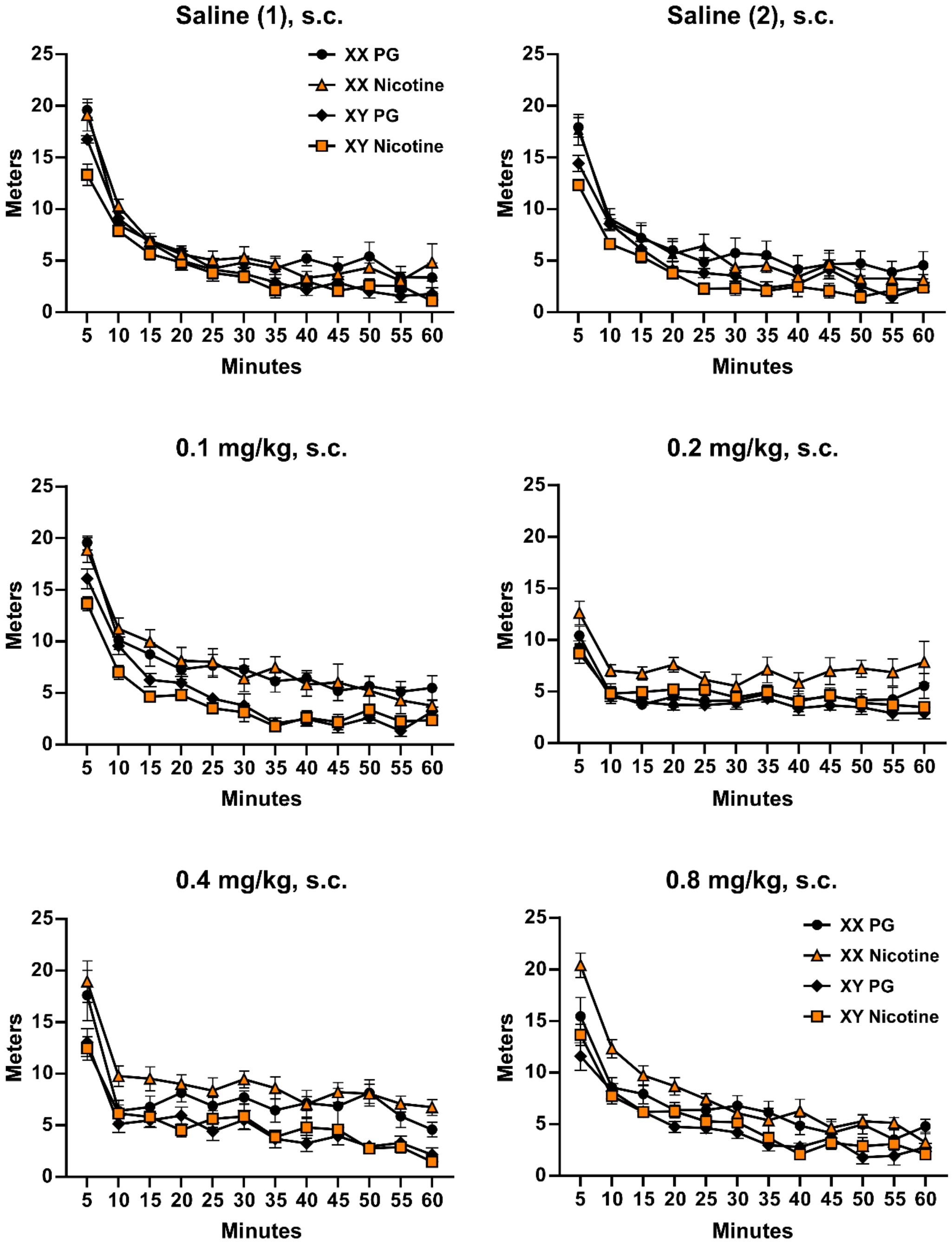


***Figure S4:*** *Mean (N=8 per group; ±SEM) distance traveled in the open arena by 5 minute time bin segments for each of six treatment conditions.*

influence of adolescent treatment was observed in the 0.4 mg/kg [Time: F (11, 308) = 64.04; P<0.0001; Sex: F (1, 28) = 22.98; P<0.0001; Interaction of Time with Sex: F (11, 308) = 1.85; P<0.05] study, however the locomotor effects of the 0.8 mg/kg dose were modulated by adolescent treatment [Time: F (11, 308) = 106.7; P<0.0001; Sex: F (1, 28) = 20.81; P<0.0001; Interaction of Time with Sex: F (11, 308) = 2.70; P<0.005; Interaction of Time with Adolescent treatment: F (11, 308) = 3.16; P<0.001; Interaction of Time, Sex and Adolescent treatment: F (11, 308) = 1.87; P<0.05].

To generalize, the rats exposed to nicotine vapor as adolescents were more resistant to the locomotor suppressing effects observed early in the session and exhibited a small locomotor stimulant effect later in the sessions and this was most pronounced in the female rats. (See **Fig. 2** of the main report for analysis of the total distance traveled).

***Experiment 3: Effect of repeated adolescent nicotine inhalation on wheel activity in adult rats***

*Baseline*: Female rats engaged in more activity than did the males on the wheels in the baseline sessions, decreased their activity more across the session and, unlike the males, increased activity from session 1 to session 2 (**Figure S5**). Since there was no impact of adolescent vapor on wheel activity for either baseline session (see below), the main analysis collapsed across this factor to compare the two baseline assessments directly (**Figure S5A**). The three-way ANOVA confirmed a significant effect of Sex [F (1, 30) = 32.35; P<0.0001], of Time within session [F (3, 90) = 13.38; P<0.0001], and of the Baseline (1 vs 2) [F (1, 30) = 38.83; P<0.0001] on wheel activity. There was also a significant effect of the interactions of Sex with Time [F (3, 90) = 3.49; P<0.05], of Sex with Baseline [F (1, 30) = 19.46; P=0.0001] and of Time with Baseline [F (3, 90) = 3.44; P<0.05]. There were no differences confirmed for the male group between baseline 1 and baseline 2 at any time during the session. The post-hoc test confirmed that female activity was significantly higher in baseline 2, compared with baseline 1, for the first 30 minutes of the

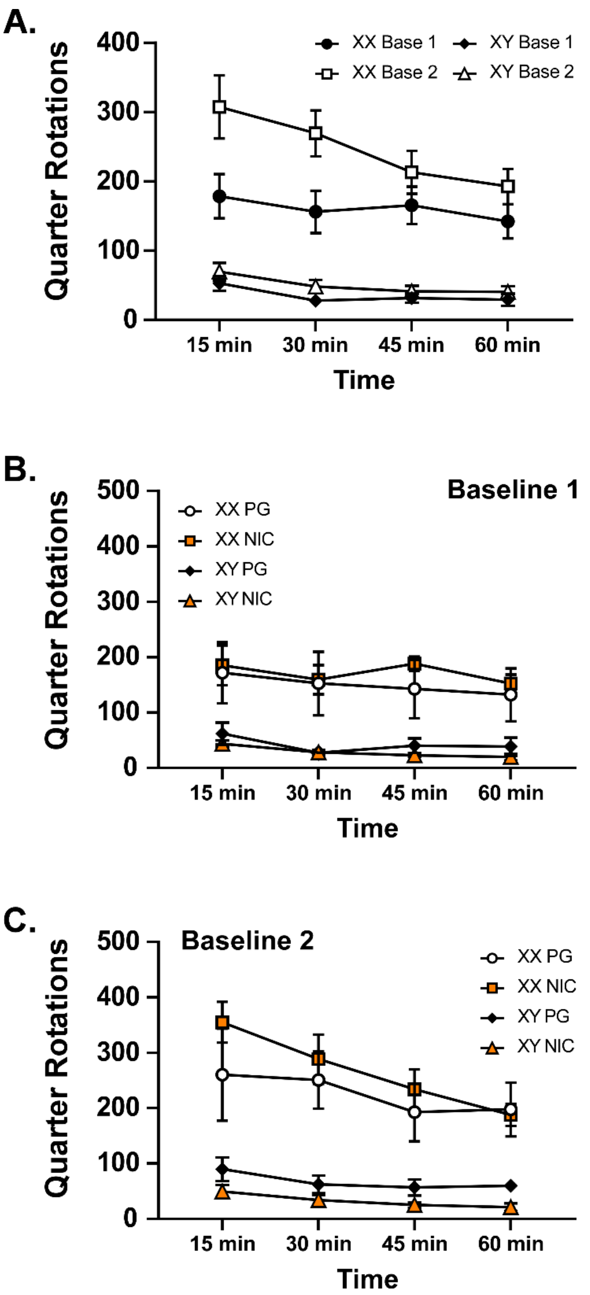


***Figure S5:*** *Mean (N=16 per group; ±SEM) wheel activity in the first two 1 h Baseline sessions by 15 minute segments for female (XX) and male (XY) rats.*

session. Female activity was significantly higher in the first 15 minutes of Baseline 2 compared with the 45- and 60-minute intervals; activity also differed between the 30 min and 60 min intervals. Activity did not significantly differ across the session for females in Baseline 1 or for males in either Baseline. Analyzed within each Baseline assessment, the activity of male and female rats was significantly different at each timepoint, except the 60-minute time for Baseline 1. In the first session (**Figure S5B**) the three way ANOVA confirmed a significant effect of Sex F (1, 28) = 22.24; P<0.0001, but not of Time, Adolescent Vapor condition or any interaction of factors. In the second baseline assessment (**Figure S5C**), the three way ANOVA confirmed a significant effect of Sex [F (1, 28) = 35.65; P<0.0001], of Time [F (3, 84) = 13.46; P<0.0001] and of the interaction of Time with Sex [F (3, 84) = 5.18; P<0.005].

*Nicotine injection*: Female rats engaged in more activity on the wheels in the s.c. injection sessions, but there was no differential effect of adolescent treatment condition on either the sex differences in wheel activity or on the effects of nicotine injection on wheel activity (**Figure S6**). The analysis confirmed significant effects of Sex, Time and the interaction of these factors in the three way ANOVAs for Saline [Time: F (1.576, 44.13) = 50.55; P<0.0001; Sex: F (1, 28) = 40.53; P<0.0001; Time x Sex Interaction: F (3, 84) = 21.67; P<0.0001], 0.2 mg/kg [Time: F (2.464, 68.99) = 22.76; P<0.0001; Sex: F (1, 28) = 45.27; P<0.0001; Time x Sex Interaction: F (3, 84) = 14.71; P<0.0001] and 0.4 mg/kg [Time: F (1.312, 36.74) = 4.30; P<0.05; Sex: F (1, 28) = 40.71; P<0.0001; Time x Sex Interaction: F (3, 84) =3.82; P<0.05] doses. There was only a significant effect of sex for the 0.8 mg/kg dose [Time: n.s.; Sex: F (1, 28) = 31.38, P<0.0001; Time x Sex Interaction: n.s.]. There were no significant effects of Adolescent Vapor condition either alone or in interaction with any other factor confirmed for any dose.

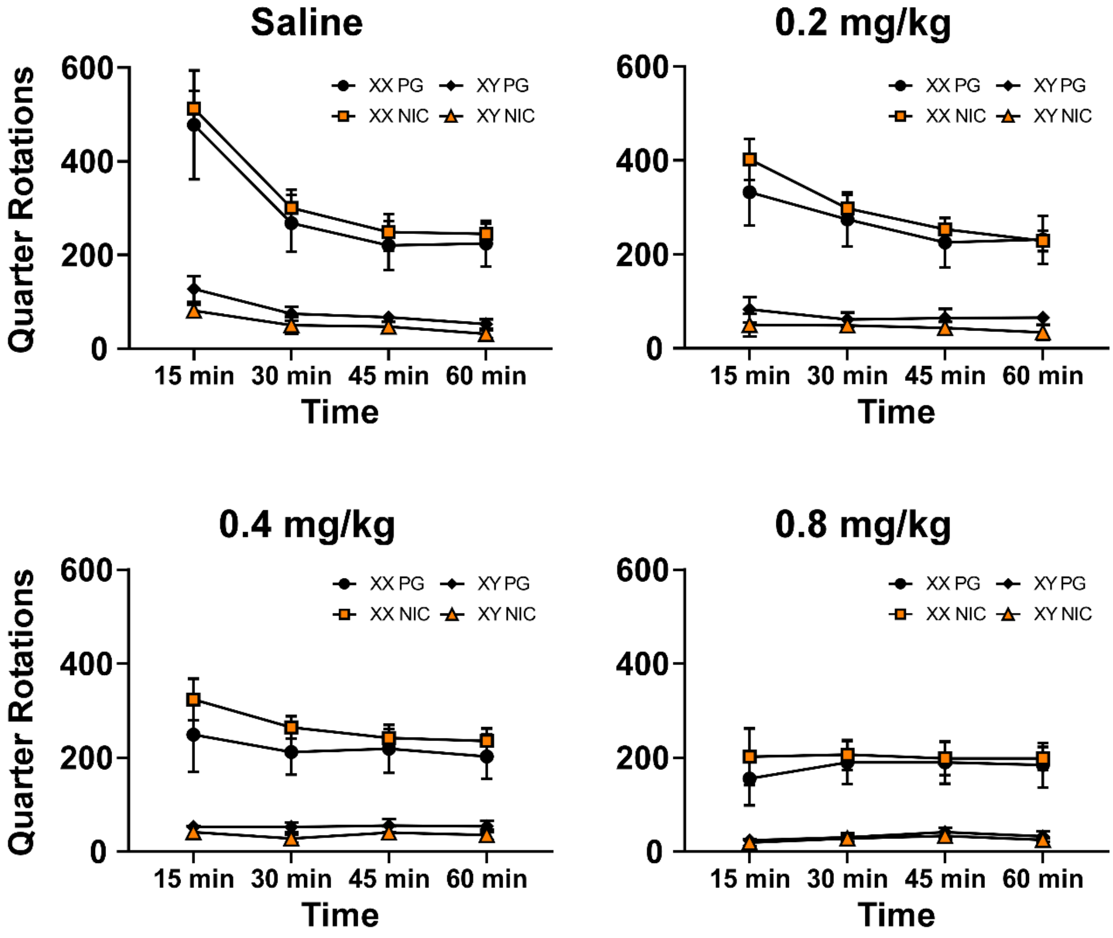


***Figure S6:*** *Mean (N=8 per group; ±SEM) wheel activity in the nicotine injection (s.c.) sessions by 15 minute segments for female (XX) and male (XY) rats exposed as adolescents to vapor from the PG vehicle or NICotine.*

#### *Experiment 4: Effect of repeated adolescent nicotine inhalation on nicotine vapor self-administration in adult rats*

**Drug-Seeking Behavior:**

Significantly more responses were made by the female group during the time-out on the drug-associated manipulandum [main effect of Concentration: $F(3, 81) = 3.13$; $P<0.05$, Sex: $F(1, 27) = 12.15$; $P<0.005$ ] when only PG vehicle vapor was available, compared with the 60 mg/mL condition, (**Figure S7A**) and on the alternate manipulandum [main effect of Concentration: $F(3, 81) = 3.99$; $P<0.05$, Sex: $F(1, 27) = 6.61$; $P<0.05$, Concentration by Sex interaction: $F(3, 81) = 4.21$; $P<0.01$ ] when only PG vehicle vapor was available, compared with all three nicotine concentrations (**Figure S7B**).

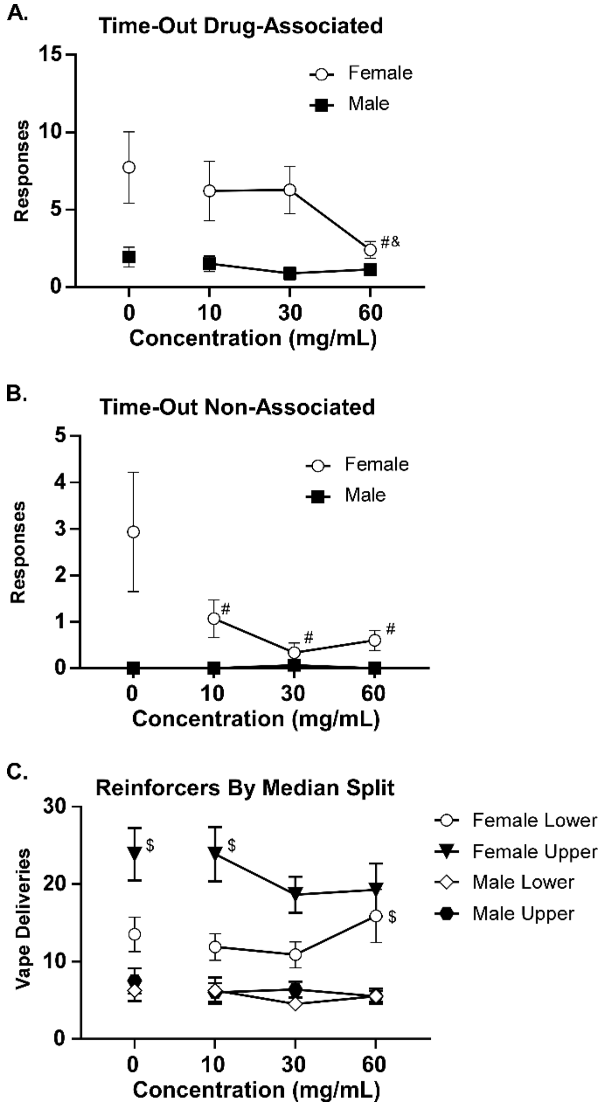


***Figure S7:*** *Mean (±SEM) responses during the post-reinforcer time-out on the A) drug-associated manipulandum and B) the alternate manipulandum for female (N=15; one outlier excluded) and male (N=16) rats collapsed across adolescent treatment condition. A significant difference from the 0 concentration within group is indicated with # C) Mean (SEM) reinforcers acquired by median split of male and female groups. A significant difference from the 30 mg/mL concentration is indicated with $.*

**Median Split Analysis:**

Significant individual differences can emerge in self-administration behavior, particularly for weaker reinforcers (**Figure S7C**). This may be due to a true lack of reinforcer efficacy for a given drug, individual differences in preferred satiety dose or even differences in preference for a given per-infusion dose. To investigate this we performed a median split on the male and female groups (regardless of adolescent treatment) based on their intake during the first 8 sessions of acquisition. The three-factor ANOVA confirmed significant effects of Concentration [$F(3, 84) = 2.9$; $P<0.05$], of Sex [$F(1, 28) = 37.16$; $P<0.0001$], of Median Split [$F(1, 28) = 6.09$; $P<0.05$] and of the interaction of Sex and Median Split [$F(1, 28) = 4.32$; $P<0.05$] on vapor deliveries obtained. The post-hoc Dunnett analysis of the Concentration factor after a two-factor follow-up ANOVA of Concentration ($F(3, 84) = 2.9$; $P<0.05$) and all four Groups ($F(3, 28) = 15.85$; $P<0.0001$) confirmed a significant difference between the 30 and 60 mg/mL concentrations for the Lower half of the female group and between the 30 and each of the 0 and 10 mg/mL concentrations for the upper half of the female group.

**FR increment:**

As described in the main report an unexpected facilities emergency disrupted behavioral testing after the 3rd FR5 session. Animals were idled for three weeks and then re-started. The analysis of the second part of the experiment contrasted the last FR1 session with the final FR5 sessions (conducted after the interruption) and then a return to FR1 for four sessions. The analysis by 3-way ANOVA confirmed that significant differences in vapor deliveries were associated with of Sex [F (1, 27) = 43.94; P<0.0001], Session [F (7, 189) = 29.47; P<0.0001], and the interaction of Session with Sex [F (7, 189) = 10.21; P<0.0001] (**Figure S8A**). There were no significant effects of Adolescent treatment alone or in interaction with other factors. Likewise, responses on the drug-associated manipulandum (**Figure S8B**) were significantly affected by Sex [F (1, 27) = 32.68; P<0.0001], Session [F (7, 189) = 18.68; P<0.0001], and the interaction of Session with Sex [F (7, 189) = 4.32; P<0.0005], as confirmed in the 3-way ANOVA. There were no significant effects of Adolescent treatment alone or in interaction with other factors on responses. There were no significant differences in percent of responses directed at the drug-associated manipulandum confirmed in the analysis (**Figure S8C**).

**PG Vapor Self-Administration / Extinction**

In one hour sessions, e.g., twice as long as for the nicotine self-administration groups, male and female rats initially obtained 11-14 vapor deliveries, on average (**Figure S9A**). As the sessions continued and the FR was incremented, vape deliveries were gradually decreased to about 5 per session. The number of responses on the vape-associated manipulandum averaged around 40 from the FR2/FR3 to FR5 stages (**Figure S9B**). The percent of responses directed at the vape-associated manipulandum increased slowly in the male rats to an average about 60% whereas the female rats' discrimination reached 80% in the early stages of the FR5 contingency (**Figure S9C**).

### Supplemental Discussion

*ENDS Delivery of Nicotine in Adolescent Rats*

The Electronic Nicotine Delivery Systems (ENDS) inhalation model is as effective in adolescent rats as it is in adult rats since nicotine inhalation reduced adolescent rats' body temperature (**Supplementary Figure S2**) in a manner similar to effects found in adult rats

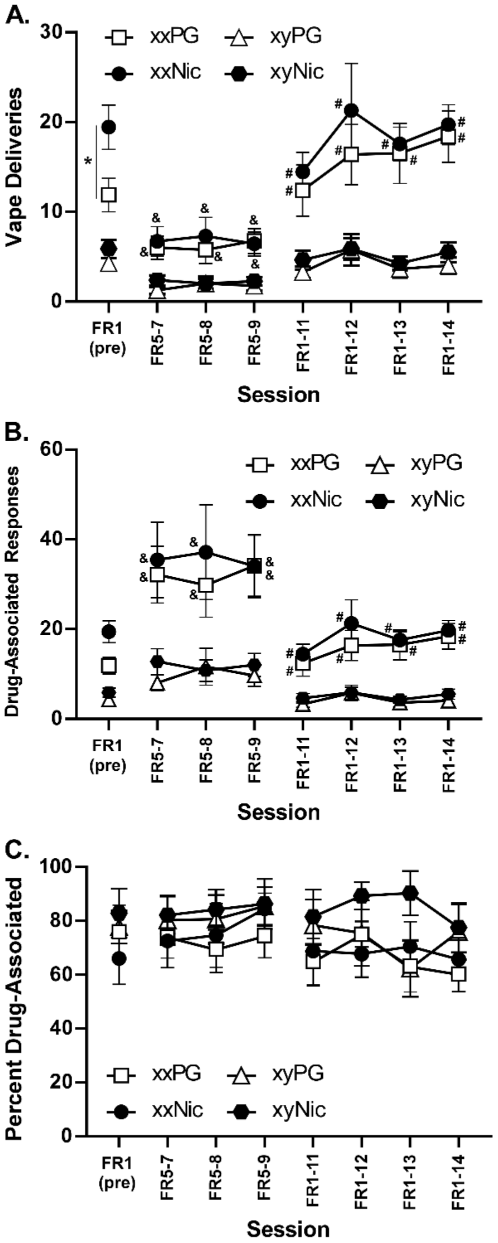


**Figure S8:** *Mean (N=7 xxNIC, others N=8 per group; ±SEM) A) nicotine (30 mg/mL) vapor deliveries, B) responses on the drug-associated manipulandum and C) percent responses on the drug-associated manipulandum obtained under Fixed Ratio (FR) 1, FR 5 and then FR1 conditions adult male (xy) and female (xx) rats exposed repeatedly during adolescence to vapor from the PG vehicle or Nicotine (30 mg/mL). A significant difference between treatment groups for a given session is indicated with *, a significant difference from the FR1 session within group with &, and a significant difference from the first FR5 session within group by #.*

(Javadi-Paydar, Mehrak et al., 2019), using similar exposure parameters (30 mg/mL nicotine, 30-minutes). Effects lasted about 20-30 minutes after the cessation of exposure, as with adult rats (Javadi-Paydar, Mehrak et al., 2019), thus, physiologically relevant conditions were used for the repeated adolescent dosing. Although we previously reported *increased* locomotor activity by telemetry after nicotine vapor inhalation (Javadi-Paydar, Mehrak et al., 2019), this was not observed in the adolescents. In those reports, however, the activity was measured within the vapor inhalation chambers without transfer to a different recording chamber, because unpublished studies found that a boost in activity associated with transfer obscured drug-related changes. The current method also produced a consistent increase in activity in all groups upon return to the recording chamber. This may be related to the increased body temperature observed in the females after PG inhalation.

Nicotine and cotinine levels after a 30-minute inhalation were consistent with those we reported for Sprague-Dawley male rats (Javadi-Paydar, Mehrak et al., 2019), but extend those data to show the time-course for conversion of nicotine to cotinine over four hours in two strains and both sexes. Although statistically reliable group differences in nicotine and cotinine were confirmed, there were only minor strain contributions in terms of mean concentrations of either cotinine or nicotine. Nicotine and cotinine levels were also similar to levels in male Wistar rats exposed to nicotine vapor (40 mg/mL nicotine concentration over 60 minutes) using slightly different puffing parameters (Montanari et al., 2020), but were higher than those reported for pregnant rats exposed to nicotine (36 mg/mL) vapor, also using different puffing parameters for 30 minutes (Breit et al., 2022). Cotinine levels were similar to those reported in a mouse model of vapor inhalation (Cooper, Akers and Henderson, 2021). The available data to date therefore show that the vapor inhalation approach is repeatable and generalizable to produced target levels of nicotine in rats.

*Sex Differences in Nicotine Self-Administration*

Significant sex differences were observed throughout the self-administration study, in both PG- and nicotine-exposed groups. Prior evidence for sex-differences in nicotine self-administration is mixed, with some studies finding no sex difference and some reporting a

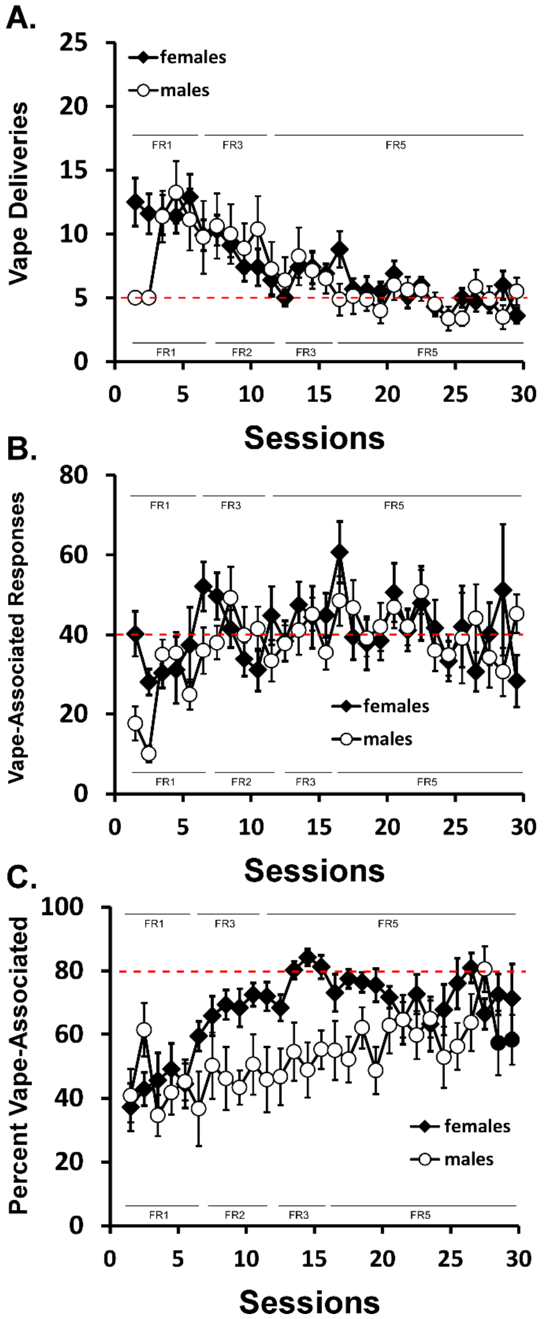


**Figure S9:** *Mean (±SEM) A) PG vapor deliveries, B) responses on the drug-associated manipulandum and C) percent responses on the drug-associated manipulandum obtained under varous Fixed Ratio (FR) conditions adult male (N=8) and female (N=10) rats. The red dotted lines are included merely to guide the eye in reference to the scale values.*

difference. The reported differences may depend on any of several methodological variables, the influences of which are poorly understood at present. A recent report found that female Sprague-Dawley rats intravenously self-administer fewer nicotine infusions than do male Sprague-Dawley rats, while no sex-difference in Long-Evans rats was observed (Leyrer-Jackson et al., 2021). There is also varied evidence of female Sprague-Dawley rats self-administering/acquiring more/faster (Sanchez et al., 2014), less/slower (Levin et al., 2011) or the same (Chaudhri et al., 2005; Donny et al., 2000; Pittenger et al., 2016) as male rats of the same strain. Similarly, female Wistar rats have been reported as self-administering/acquiring more/faster (Chellian et al., 2021; Uribe et al., 2020), less/slower (Swalve, Smethells and Carroll, 2016) or the same as male rats of the same strain and a similar diversity of sex-associated outcome has been reported for Long Evans rats (Leyrer-Jackson et al., 2021; Li et al., 2014). A study of heterogeneous stock adolescent rats derived from eight founder strains reported increased oral nicotine intake in female rats (Wang et al., 2018). Another study found that female rats intravenously self-administer higher amounts of nicotine only under FR5, but not FR1, and as in the present study, responses only partially compensated (i.e.12-19 responses under FR1, 30-70 under FR5) for the increased response requirement (Chaudhri et al., 2005). Male and female Sprague-Dawley rats intravenously self-administer identical numbers of infusions of nicotine when self-administration is started in adolescence (PND 34) after food-reinforced operant training (Carreno and Lotfipour, 2023) and adult female and male Wistar rats self-administered similar amounts of *nicotine vapor*, also following food-reinforced operant training (Lallai et al., 2021). Given this diversity of outcomes it is clear that the variation in methodological approach contributes strongly to the appearance, or absence, of sex differences in rat nicotine self-administration.

*Individual Differences in Female Rat Nicotine Self-Administration*

Despite the fact that self-administration paradigms are designed to let individual animals titrate their overall session intake, and to some extent the rate of that intake, traditional group-based procedures and analysis may sometimes limit the ability to infer drug reinforcement in some individuals or across groups. For example while 60% of rats acquired the self-administration of 3,4-methylenedioxymethamphetamine (MDMA) in roughly 10 session compared to 100% that acquired cocaine self-administration in one study (Schenk et al., 2007), the same group showed that many “non-acquiring” self-administration of MDMA will meet acquisition criteria after some 20-25 sessions (Schenk et al., 2012). Likewise, the impact of an anti-oxycodone vaccine differed across more-preferring and less-preferring halves of the distribution (Nguyen et al., 2018). It is also the case that there can be individual left/right shifts of the curve generated in a dose-substitution procedure, that produce a group mean of no significant dose difference. In the present data, the more preferring Upper half female rats and the less preferring Lower half female rats exhibited seeming dose preference differences. The latter responded for more vapor deliveries when the concentration was increased above the training concentration. This suggests that one potential reason for them being in the less preferring half during acquisition is that the training dose was below that necessary for them to express full drug preference. The Upper half females, on the other hand, engaged in increased seeking behavior when the concentration was lowered below the training (30 mg/mL) concentration.

**PG Vapor Self-Administration / Extinction**

Rats’ initial self-administration of PG vapor was about 11-14 vapor deliveries in one hour sessions, e.g., twice as long as the sessions for the nicotine groups. The female rats in the main study initially obtained an average of 20 vape deliveries in 30 minute sessions and finished acquisition between 12 (adolescent PG) or 20 (adolescent nicotine) vape deliveries. This represents approximately 2-4 times the rate of PG self-administration expressed by the females. The males’ self-administration of PG was similar to the female rats across all phases (**Figure S9A**), in striking contrast to the sex difference in the nicotine self-administration

(**Figure 3**). The male nicotine groups initially obtained about 10 vape deliveries but eventually declined to around 5 or 6 at the end of acquisition, that is, from 2x to 1x the *rate* of the male PG self-administration. As the sessions continued and the FR was incremented, mean PG vape deliveries were eventually decreased from 10 to about 5 per session in both male and female groups. Compared with the female rats in the main study this is about a quarter of the behavioral rate at the third FR5 session (**Figure 4**), while recovery was occurring, and about half of the rate observed at the end of the FR5 sequence (**Figure S8**). The fact that the stable response rate for vehicle vapor is about 5 deliveries per hour under an FR5 contingency for female rats after extensive experience, whereas the nicotine rate is 10 deliveries per hour under FR5 is further support for the inference that the female rats in the main study were seeking nicotine, i.e., self-administering.

### Supplemental Literature Cited


Breit, K.R., Rodriguez, C.G., Hussain, S., Thomas, K.J., Zeigler, M., Gerasimidis, I., Thomas, J.D., 2022. A Model of Combined Exposure to Nicotine and Tetrahydrocannabinol via Electronic Cigarettes in Pregnant Rats. Front Neurosci 16, 866722, doi: 10.3389/fnins.2022.866722.

Carreno, D., Lotfipour, S., 2023. Male and Female Sprague Dawley Rats Exhibit Equivalent Natural Reward, Nicotine Self-Administration, Extinction, and Reinstatement During Adolescent-Initiated Behaviors. Nicotine & tobacco research : official journal of the Society for Research on Nicotine and Tobacco 25(5), 1039-1046, doi: 10.1093/ntr/ntac234.

Chaudhri, N., Caggiula, A.R., Donny, E.C., Booth, S., Gharib, M.A., Craven, L.A., Allen, S.S., Sved, A.F., Perkins, K.A., 2005. Sex differences in the contribution of nicotine and nonpharmacological stimuli to nicotine self-administration in rats. Psychopharmacology 180(2), 258-266, doi: 10.1007/s00213-005-2152-3.

Chellian, R., Behnood-Rod, A., Wilson, R., Febo, M., Bruijnzeel, A.W., 2021. Adolescent nicotine treatment causes robust locomotor sensitization during adolescence but impedes the spontaneous acquisition of nicotine intake in adult female Wistar rats. Pharmacology, biochemistry, and behavior 207, 173224, doi: 10.1016/j.pbb.2021.173224.

Cooper, S.Y., Akers, A.T., Henderson, B.J., 2021. Flavors Enhance Nicotine Vapor Self-administration in Male Mice. Nicotine & tobacco research : official journal of the Society for Research on Nicotine and Tobacco 23(3), 566-572, doi: 10.1093/ntr/ntaa165.

Donny, E.C., Caggiula, A.R., Rowell, P.P., Gharib, M.A., Maldovan, V., Booth, S., Mielke, M.M., Hoffman, A., McCallum, S., 2000. Nicotine self-administration in rats: estrous cycle effects, sex differences and nicotinic receptor binding. Psychopharmacology 151(4), 392-405, doi:

Garber, J.C., Barbee, R.W., Bielitzki, J.T., Clayton, L.A., Donovan, J.C., Hendriksen, C.F.M., Kohn, D.F., Lipman, N.S., Locke, P.A., Melcher, J., Quimby, F.W., Turner, P.V., Wood, G.A., Wurbel, H., 2011. Guide for the Care and Use of Laboratory Animals, 8th Edition. National Academies Press, Washington D.C.

Gilpin, N.W., Wright, M.J., Jr., Dickinson, G., Vandewater, S.A., Price, J.U., Taffe, M.A., 2011. Influences of activity wheel access on the body temperature response to MDMA and methamphetamine. Pharmacology, biochemistry, and behavior 99(3), 295-300, doi: 10.1016/j.pbb.2011.05.006.

Gutierrez, A., Creehan, K.M., Taffe, M.A., 2021. A vapor exposure method for delivering heroin alters nociception, body temperature and spontaneous activity in female and male rats. J Neurosci Methods 348, 108993, doi: 10.1016/j.jneumeth.2020.108993.

Javadi-Paydar, M., Creehan, K.M., Kerr, T.M., Taffe, M.A., 2019a. Vapor inhalation of cannabidiol (CBD) in rats. Pharmacology, biochemistry, and behavior 184, 172741, doi: 10.1016/j.pbb.2019.172741.

Javadi-Paydar, M., Kerr, T.M., Harvey, E.L., Cole, M., Taffe, M.A., 2019. Effects of Nicotine and THC Vapor Inhalation Administered by An Electronic Nicotine Delivery System (ENDS) in Male Rats. Drug & Alcohol Dependence in press, doi:

Javadi-Paydar, M., Kerr, T.M., Harvey, E.L., Cole, M., Taffe, M.A., 2019b. Effects of nicotine and THC vapor inhalation administered by an electronic nicotine delivery system (ENDS) in male rats. Drug and alcohol dependence 198, 54-62, doi: 10.1016/j.drugalcdep.2019.01.027.

Lallai, V., Chen, Y.C., Roybal, M.M., Kotha, E.R., Fowler, J.P., Staben, A., Cortez, A., Fowler, C.D., 2021. Nicotine e-cigarette vapor inhalation and self-administration in a rodent model: Sex- and nicotine delivery-specific effects on metabolism and behavior. Addict Biol 26(6), e13024, doi: 10.1111/adb.13024.

Levin, E.D., Slade, S., Wells, C., Cauley, M., Petro, A., Vendittelli, A., Johnson, M., Williams, P., Horton, K., Rezvani, A.H., 2011. Threshold of adulthood for the onset of nicotine self-

administration in male and female rats. Behavioural brain research 225(2), 473-481, doi: 10.1016/j.bbr.2011.08.005.
Leyrer-Jackson, J.M., Overby, P.F., Bull, A., Marusich, J.A., Gipson, C.D., 2021. Strain and sex matters: Differences in nicotine self-administration between outbred and recombinase-driver transgenic rat lines. Experimental and clinical psychopharmacology 29(4), 375-384, doi: 10.1037/pha0000376.
Li, S., Zou, S., Coen, K., Funk, D., Shram, M.J., Le, A.D., 2014. Sex differences in yohimbine-induced increases in the reinforcing efficacy of nicotine in adolescent rats. Addict Biol 19(2), 156-164, doi: 10.1111/j.1369-1600.2012.00473.x.
Miller, M.L., Moreno, A.Y., Aarde, S.M., Creehan, K.M., Vandewater, S.A., Vaillancourt, B.D., Wright, M.J., Jr., Janda, K.D., Taffe, M.A., 2013. A methamphetamine vaccine attenuates methamphetamine-induced disruptions in thermoregulation and activity in rats. Biol Psychiatry 73(8), 721-728, doi: 10.1016/j.biopsych.2012.09.010.
Montanari, C., Kelley, L.K., Kerr, T.M., Cole, M., Gilpin, N.W., 2020. Nicotine e-cigarette vapor inhalation effects on nicotine & cotinine plasma levels and somatic withdrawal signs in adult male Wistar rats. Psychopharmacology 237(3), 613-625, doi: 10.1007/s00213-019-05400-2.
Nguyen, J.D., Aarde, S.M., Vandewater, S.A., Grant, Y., Stouffer, D.G., Parsons, L.H., Cole, M., Taffe, M.A., 2016. Inhaled delivery of Delta(9)-tetrahydrocannabinol (THC) to rats by e-cigarette vapor technology. Neuropharmacology 109, 112-120, doi: 10.1016/j.neuropharm.2016.05.021.
Nguyen, J.D., Creehan, K.M., Kerr, T.M., Taffe, M.A., 2020. Lasting effects of repeated Δ(9) - tetrahydrocannabinol vapour inhalation during adolescence in male and female rats. Br J Pharmacol 177(1), 188-203, doi: 10.1111/bph.14856.
Nguyen, J.D., Grant, Y., Creehan, K.M., Hwang, C.S., Vandewater, S.A., Janda, K.D., Cole, M., Taffe, M.A., 2019. Delta(9)-tetrahydrocannabinol attenuates oxycodone self-administration under extended access conditions. Neuropharmacology 151, 127-135, doi: 10.1016/j.neuropharm.2019.04.010.
Nguyen, J.D., Hwang, C.S., Grant, Y., Janda, K.D., Taffe, M.A., 2018. Prophylactic vaccination protects against the development of oxycodone self-administration. Neuropharmacology 138, 292-303, doi: 10.1016/j.neuropharm.2018.06.026.
Pittenger, S.T., Swalve, N., Chou, S., Smith, M.D., Hoonakker, A.J., Pudiak, C.M., Fleckenstein, A.E., Hanson, G.R., Bevins, R.A., 2016. Sex differences in neurotensin and substance P following nicotine self-administration in rats. Synapse (New York, N.Y.) 70(8), 336-346, doi: 10.1002/syn.21907.
Sanchez, V., Moore, C.F., Brunzell, D.H., Lynch, W.J., 2014. Sex differences in the effect of wheel running on subsequent nicotine-seeking in a rat adolescent-onset self-administration model. Psychopharmacology 231(8), 1753-1762, doi: 10.1007/s00213-013-3359-3.
Schenk, S., Colussi-Mas, J., Do, J., Bird, J., 2012. Profile of MDMA Self-Administration from a Large Cohort of Rats: MDMA Develops a Profile of Dependence with Extended Testing. Journal of Drug and Alcohol Research 1(1), 1-6, doi: doi:10.4303/jdar/235602.
Schenk, S., Hely, L., Lake, B., Daniela, E., Gittings, D., Mash, D.C., 2007. MDMA self-administration in rats: acquisition, progressive ratio responding and serotonin transporter binding. Eur J Neurosci 26(11), 3229-3236, doi: 10.1111/j.1460-9568.2007.05932.x.
Swalve, N., Smethells, J.R., Carroll, M.E., 2016. Sex differences in the acquisition and maintenance of cocaine and nicotine self-administration in rats. Psychopharmacology 233(6), 1005-1013, doi: 10.1007/s00213-015-4183-8.
Taffe, M.A., Creehan, K.M., Vandewater, S.A., 2015. Cannabidiol fails to reverse hypothermia or locomotor suppression induced by Delta(9) - tetrahydrocannabinol in Sprague-Dawley rats. British journal of pharmacology 172(7), 1783-1791, doi: 10.1111/bph.13024.
Taffe, M.A., Creehan, K.M., Vandewater, S.A., Kerr, T.M., Cole, M., 2021. Effects of Delta(9)-tetrahydrocannabinol (THC) vapor inhalation in Sprague-Dawley and Wistar rats. Experimental and clinical psychopharmacology 29(1), 1-13, doi: 10.1037/pha0000373.
Uribe, K.P., Correa, V.L., Pinales, B.E., Flores, R.J., Cruz, B., Shan, Z., Bruijnzeel, A.W., Khan, A.M., O'Dell, L.E., 2020. Overexpression of corticotropin-releasing factor in the nucleus accumbens enhances the reinforcing effects of nicotine in intact female versus male and ovariectomized female rats. Neuropsychopharmacology : official publication of the American College of Neuropsychopharmacology 45(2), 394-403, doi: 10.1038/s41386-019-0543-0.
Wang, T., Han, W., Chitre, A.S., Polesskaya, O., Solberg Woods, L.C., Palmer, A.A., Chen, H., 2018. Social and anxiety-like behaviors contribute to nicotine self-administration in adolescent outbred rats. Sci Rep 8(1), 18069, doi: 10.1038/s41598-018-36263-w.
